# The HSPG Syndecan is a core organizer of cholinergic synapses in *C. elegans*

**DOI:** 10.1101/2020.11.25.395806

**Authors:** Xin Zhou, Camille Vachon, Mélissa Cizeron, Océane Romatif, Hannes E. Bülow, Maëlle Jospin, Jean-Louis Bessereau

**Author notes:** Co-first authors (Xin Zhou, Camille Vachon). Corresponding author Jean-Louis BESSEREAU, Institut NeuroMyoGene, Laboratory of Genetics and Neurobiology of *C. elegans*, Univ Lyon, Université Claude Bernard Lyon 1, CNRS UMR 5310, INSERM U 1217, Faculté de Médecine et de Pharmacie, 3ème étage-Aile D, 8, Avenue Rockefeller, 69008 Lyon - France.

## Abstract

The extracellular matrix has emerged as an active component of chemical synapses regulating synaptic formation, maintenance and homeostasis. The heparan sulfate proteoglycan syndecans are known to regulate cellular and axonal migration in the brain. They are also enriched at synapses, but their synaptic functions remain more elusive. Here we show that SDN-1, the sole ortholog of syndecan in *C. elegans*, is absolutely required for the synaptic clustering of homomeric α7-like N-acetylcholine receptors (AChR) and regulates the synaptic content of heteromeric L-AChRs. SDN-1 is concentrated at neuromuscular junctions (NMJs) by the neurally-secreted synaptic organizer Ce-Punctin/MADD-4, which also activates the transmembrane netrin receptor DCC. Those cooperatively recruit the FARP and CASK orthologues that localize N-AChRs at cholinergic NMJs through physical interactions. Therefore, SDN-1 stands at the core of the cholinergic synapse organization by bridging the extracellular synaptic determinants to the intracellular synaptic scaffold that controls the postsynaptic receptor content.

## INTRODUCTION

Chemical synapses are specialized cellular junctions that support directional transfer of information between excitable cells. However, glial cells, by engaging intimate contacts with neuronal components at synaptic sites, were identified as important synaptic players, leading to the notion of the ‘tripartite synapse’ (Araque et al., 1999; Halassa et al., 2007). In addition, the extracellular matrix (ECM) that fills the synaptic cleft and the perisynaptic extracellular space emerged as an active element that orchestrates synaptogenesis, synaptic maintenance and synaptic function, thus leading to the concept of the ‘tetrapartite synapse’ (Dityatev and Rusakov, 2011; Dityatev et al., 2006; Ferrer-Ferrer and Dityatev, 2018). The synaptic ECM, also referred to as ‘synaptomatrix’ (Dani and Broadie, 2012; Heikkinen et al., 2014), forms a dense and composite environment. It is made of secreted glycoproteins and proteoglycans intermingled with the extracellular regions of pre- and postsynaptic membrane proteins and outer-leaflet glycolipids. Signaling between pre- and postsynaptic partners necessarily occurs through this specific environment, which consequently impacts all aspects of synaptic biology (Heikkinen et al., 2014; Kurshan et al., 2014).

Within the synaptomatrix, some glycoproteins such as heparan sulfate proteoglycans (HSPG) fulfill the role of *bona fide* synaptic organizers (Condomitti and de Wit, 2018a; Yuzaki, 2018). HSPGs are either secreted or membrane-associated core proteins bearing covalently-linked chains of long disaccharide repeats (Sarrazin et al., 2011). Enzymatic modifications of the glycan chains, including epimerization, deacetylations and sulfation at various positions, creates highly-negatively charged chains with distinct structural motifs (Lindahl and Li, 2009). HSPGs bind multiple partners and can promote or regulate synaptogenesis (Song and Kim 2013). The founding member of this diverse group is agrin, a large HSPG secreted by motoneurons at the vertebrate neuromuscular junctions (NMJs), which is absolutely required for NMJ differentiation and postsynaptic clustering of acetylcholine receptors (AChRs) (Burden et al., 2018; Li et al., 2018; Swenarchuk, 2019). Since then, an increasing number of HSPGs have been demonstrated to function as synaptic regulators (reviewed in Condomitti and de Wit, 2018). For example, in the hippocampus, the GPI-anchored glypican GPC4 binds the postsynaptic adhesion molecule LRRTM4 (de Wit et al., 2013) or the orphan receptor GPR158 (Condomitti et al., 2018) and promotes presynaptic differentiation through LAR-family receptor protein tyrosine phosphatases (RPTP). At other synapses, GPC4 is secreted by astrocytes and binds presynaptic RPTPδ and RPTPσ to trigger the release of neuronal pentraxin 1, which in turn increases AMPAR content (Allen et al., 2012; Farhy-Tselnicker et al., 2017).

Syndecans are a class of transmembrane HSPGs also shown to play roles at synapses(Saied-Santiago and Bülow, 2018). In Drosophila, syndecan localizes to glutamatergic NMJs. It competes with the glypican Dallylike for binding to the LAR-family receptor protein tyrosine phosphatase dLAR and promotes the growth of presynaptic terminals (Johnson et al., 2006). Although syndecan was initially reported to be acting cell-autonomously in the motoneuron terminal, further evidence suggests that it might be provided post-synaptically by muscle cells (Nguyen et al., 2016). In mammals, the syndecan family has 4 members. Syndecan 2 is expressed in neurons of young and adult rats (Hsueh and Sheng, 1999) and progressively concentrates at asymmetric synapses in most regions of the adult rat brain (Hsueh et al., 1998). It can promote the formation of dendritic spines in cell culture (Ethell and Yamaguchi, 1999; Ethell et al., 2000; Lin et al., 2007), notably by facilitating FGF22 targeting to dendritic filopodia and spines (Hu et al., 2016). In turn, FGF22 promotes presynaptic differentiation hence coordinating pre- and postsynaptic maturation in cultured neurons (Hu et al., 2016). *Sdc2*^*−/−*^ mice are born alive with angiogenic defects, but the characterization of their nervous system has not been reported (Corti et al., 2019). Syndecan-3 is also highly expressed in the developing and adult brain, but is almost exclusively detected in axonal tracts (Hsueh and Sheng, 1999). Inactivation of *Sdc3* does not cause global abnormalities of the brain anatomy (Kaksonen et al., 2002) but the laminar structure of the cerebral cortex is perturbed as a result of impaired radial migration (Hienola et al., 2006). No modification of synaptic density nor basal transmission were reported in the hippocampus. However, *Sdc3*^*−/−*^ mice have enhanced LTP in the CA1 area and impaired hippocampus-dependent memory behaviors (Kaksonen et al., 2002), suggesting that syndecan 3 is required for synaptic plasticity.

HSPGs, in particular SDN-1, the sole syndecan member in *C. elegans*, is expressed and functions in the nervous system (Minniti et al., 2004; Rhiner et al., 2005). Moreover, indirect evidence suggests that heparan sulfates are highly enriched at *C. elegans* NMJs and are carried by SDN-1 (Attreed et al., 2012). We therefore used the *C. elegans* NMJ as a genetically-tractable model to investigate further the synaptic function of SDN-1. In *C. elegans* each body-wall muscle cell receives excitatory and inhibitory innervation from cholinergic and GABAergic motoneurons. The cholinergic versus GABAergic identity of postsynaptic domains is specified by the anterograde synaptic organizer Ce-Punctin/MADD-4 (Pinan-Lucarré et al., 2014). This evolutionarily conserved ECM protein belongs to the poorly characterized ADAMTS-like proteins that contain multiple thrombospondin type-1 repeat (TSR) and immunoglobulin (Ig) domains as well as structurally-unsolved domains in common with the ADAMTS family (Apte, 2009). The functions of its vertebrate orthologs *Punctin1/ADAMTSL1* and *Punctin2/ADAMTSL3* are ill-defined, but *Punctin2* is expressed in the brain and was identified as a susceptibility gene for schizophrenia (Dow et al., 2011).

*Ce-punctin* generates long (L) and short (S) isoforms by use of alternative promoters. The combination of isoforms present in the synaptic cleft controls the identity of the postsynaptic domains. Punctin L is exclusively secreted by cholinergic neurons and triggers the postsynaptic localization of two types of ionotropic acetylcholine receptors (AChRs): levamisole-sensitive AChRs (L-AChRs) that are heteromeric and can be activated by the nematode-specific cholinergic agonist levamisole, and the nicotine-sensitive AChRs (N-AChRs) that are homomeric, activated by nicotine, and evolutionarily very close to the α7 AChRs expressed in the mammalian brain. L-AChRs interact with an extracellular scaffold (Gally et al., 2004; Gendrel et al., 2009; Rapti et al., 2011) to form microclusters that are positioned at synapses by Punctin L. The Punctin-dependent clustering of N-AChRs relies on a distinct uncharacterized pathway. The short isoform Punctin S is secreted by both cholinergic and GABAergic motoneurons. It triggers the postsynaptic clustering of type A GABA receptors (GABA_A_Rs) at GABAergic NMJs. At cholinergic NMJs, it associates with Punctin L and inhibits the inappropriate recruitment of GABA_A_Rs by Punctin L. Punctin S controls the clustering of GABA_A_Rs by two convergent molecular pathways (for review, see Zhou and Bessereau 2019). First, Punctin S binds and clusters the synaptic adhesion molecule NLG-1/neuroligin in front of GABAergic boutons, which controls the synaptic localization of GABA_A_Rs (Maro et al., 2015; Tu et al., 2015). Second, it binds, recruits, and likely activates the netrin receptor UNC-40/DCC, which controls the synaptic content of GABA_A_Rs (Tu et al., 2015). UNC-40/DCC nucleates an intracellular scaffold by physically interacting with FRM-3, a FERM (p4.1, Ezrin, Radixin, Moesin) protein orthologous to FARP1/2 (Zhou et al., 2020). FRM-3 multimerizes and recruits LIN-2, the ortholog of CASK (Calcium calmodulin dependent Serine/Threonine kinase), which might provide a hub to physically connect the GABA_A_Rs, FRM-3/FARP and NLG-1/neuroligin. Except NLG-1, these molecules are also present at cholinergic NMJs but their precise function was not characterized.

Here we show that SDN-1/syndecan is concentrated at NMJs by Punctin and modulates the content of receptors at excitatory and inhibitory NMJs. Remarkably, SDN-1 is the core component supporting the clustering of homomeric N-AChRs. Through physical interaction with LIN-2/CASK and FRM-3/FARP, SDN-1 stabilizes a DCC-dependent synaptic scaffold at cholinergic NMJs, which in turn recruits N-AChRs. Since all these proteins are evolutionarily conserved, these results provide a framework to analyze the molecular mechanisms controlling the localization of alpha-7 AChRs in mammalian neurons.

## RESULTS

### Syndecan / SDN-1 is a synaptic protein at NMJs

Previous studies suggested that SDN-1 localizes in synapse-rich regions of the *C. elegans* nervous system based on the strong decrease of GAG immunoreactivity in the nerve ring and the nerve cords of *sdn-1* mutants (Attreed et al., 2012). To define the precise localization of the SDN-1 HSPG, we used the CRISPR/Cas9 technique to insert the fluorescent protein mNeonGreen (mNG) at the N-terminus of SDN-1 immediately after its signal peptide (Fig. 1A). In adult worms, mNG-SDN-1 was detected at high level in the nerve ring and along the ventral and dorsal cords (Fig. 1B). The fusion protein mNG-SDN-1 appeared functional based on the absence of abnormal visible phenotypes of knock-in animals, the absence of obvious axonal outgrowth defects, and the normal content of ACh and GABA_A_ receptors at NMJs as compared to the wild type (Fig. S1).

**Figure 1.**
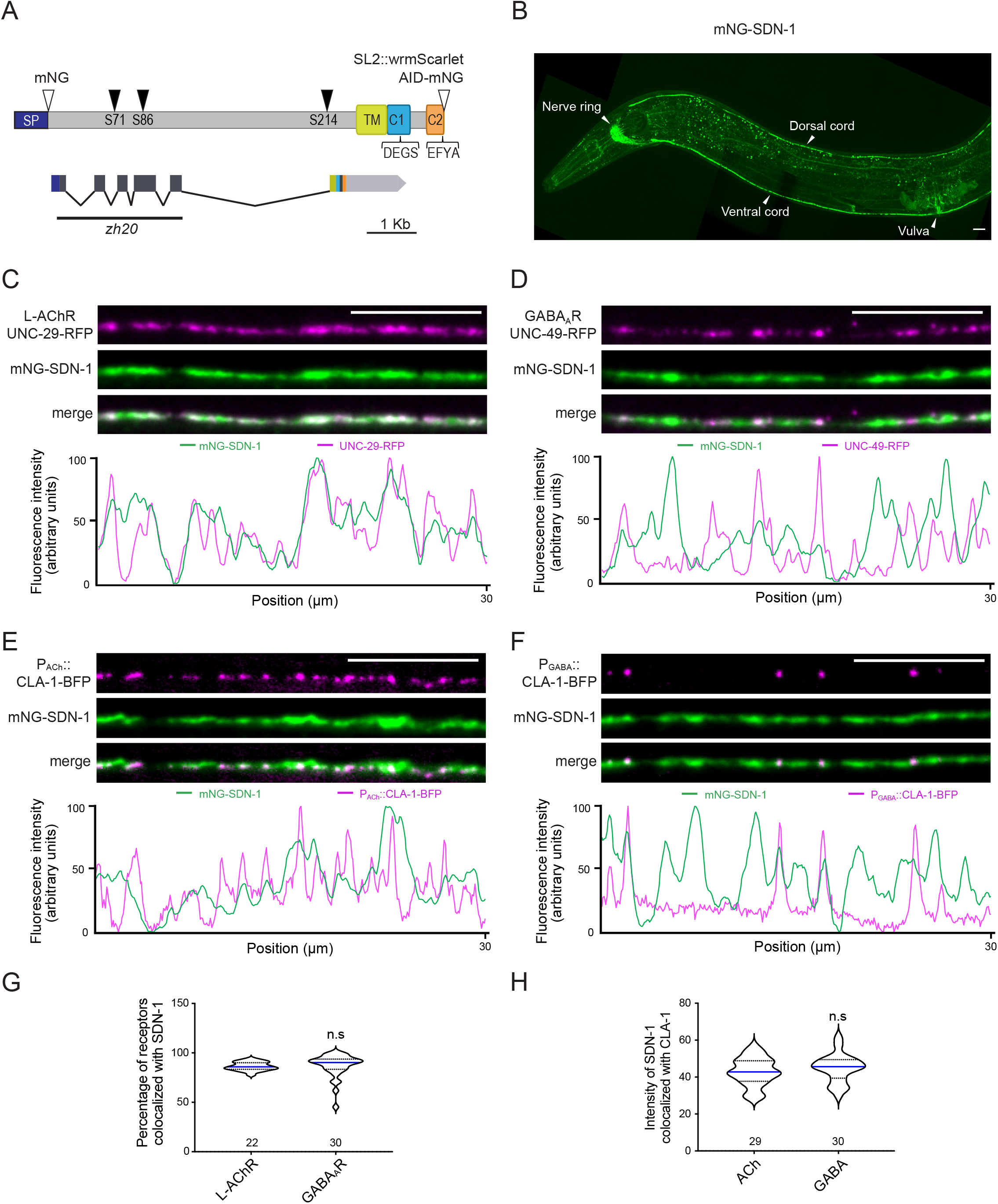
SDN-1 is enriched at NMJs. (A) Predicted structure of SDN-1 protein (288 aa) and *sdn-1* locus (2921 bp). SP (signal peptide, dark blue box), TM (transmembrane, light green box), C1 (containing the DEGS ezrin binding motif, light blue box), C2 (containing the PDZ binding motif EFYA, orange box). Three putative sites (black arrowheads) for GAG anchoring have been predicted on serine 71, 86 and 214. Open arrowheads indicate the knock-in sites of fluorescent tags: mNeonGreen (mNG) right after SP, auxin-inducible degron AID-mNG or SL2-wrmScarlet (wormScarlet) right after the PDZ binding motif. The gray bar marks the location of the 1258 bp deletion in *sdn-1(zh20)* mutant allele. Boxes represent exons. (B) High expression level of mNG-SDN-1 is detected in the nerve ring and along the nerve cords. Scale bar is 10 μm. (C-D) Confocal detection and fluorescence profiles along the dorsal cord of mNG-SDN-1 and TagRFP-T labeled receptors: L-AChR is labelled from the *unc-29-TagRFP-T* knock-in allele and GABA_A_R is labelled from the *TagRFP-T-unc-49* knock-in allele. (E-F) Confocal detection and fluorescence profiles along the dorsal cord of mNG-SDN-1 and the presynaptic active zone marker CLA-1-BFP under the control of the cholinergic neuron-specific promoter *Punc-17* (F) or under the control of the GABAergic neuron-specific promoter *Punc-47* (G). (G) Percentage of L-AChRs and GABA_A_Rs colocalized with mNG-SDN-1, calculated using Manders coefficient on processed images (see material and methods for details). (H) Mean intensity of the mNG-SDN-1 signal specifically colocalized with cholinergic or GABAergic synapses, calculated following image processing (see material and methods for details). In this figure and all the other confocal figures, pictures are sums of Z-stacks acquired along the dorsal nerve cord by a spinning disk confocal microscope; anterior is to the left, scale bar is 10 μm. In this figure and all the other figures, data distribution in each group is presented as violin plots showing lower and upper quartiles (dotted lines), median (blue line). The number of animals analyzed is indicated on the violin plots. Mann-Whitney test, n.s: not significant. See also Fig. S1.

To test if mNG-SDN-1 was enriched at synapses, we used knock-in strains expressing L-AChRs or GABARs tagged with red fluorescent proteins. Visualization of the ventral and dorsal nerve cords at high magnification showed that mNG-SDN-1 was highly concentrated at NMJs. Quantitative analysis showed that both ACh and GABA_A_ receptors almost completely colocalized with syndecan (Fig. 1C, D, G). We also measured the SDN-1 density at each synaptic type using the presynaptic marker CLA-1/Clarinet (Xuan et al., 2017), which accurately delineates active zones (Fig. 1E, F), and we observed that SDN-1 was equally abundant at excitatory and inhibitory NMJs (Fig. 1H).

### Syndecan controls the synaptic localization of α7-like N-AChRs

To identify putative functions of SDN-1 at synapses, we analyzed NMJs in *sdn-1(zh20)* null mutants, hereinafter referred to as *sdn-1(0)*. The number of synapses was not changed in *sdn-1(0)* mutants as compared to the wild type, based on presynaptic markers of either cholinergic or GABAergic terminals (Fig. 2A, S2A). By contrast, we observed strong alterations of postsynaptic receptor distribution. First, the synaptic content of heteromeric L-AChRs was decreased by roughly 60 % in *sdn-1(0)* mutants as compared to the wild type, based on the fluorescence of the UNC-29-RFP subunit expressed from a knock-in allele (Fig. 2B). The remaining receptors were still clustered at synapses. Second, homomeric N-AChRs, that are homologous to α7 nicotinic receptors in mammals, were almost undetectable based on ACR-16-wrmScarlet fluorescence (Fig. 2C). Third, synaptic GABA_A_R content was also reduced, albeit to a lesser extent, with a roughly 35 % decrease of UNC-49-RFP fluorescence at synapses (Fig. S2B). Because of the more modest impact of syndecan loss at GABAergic NMJs, we subsequently focused our analysis on SDN-1-dependent localization of AChRs.

**Figure 2.**
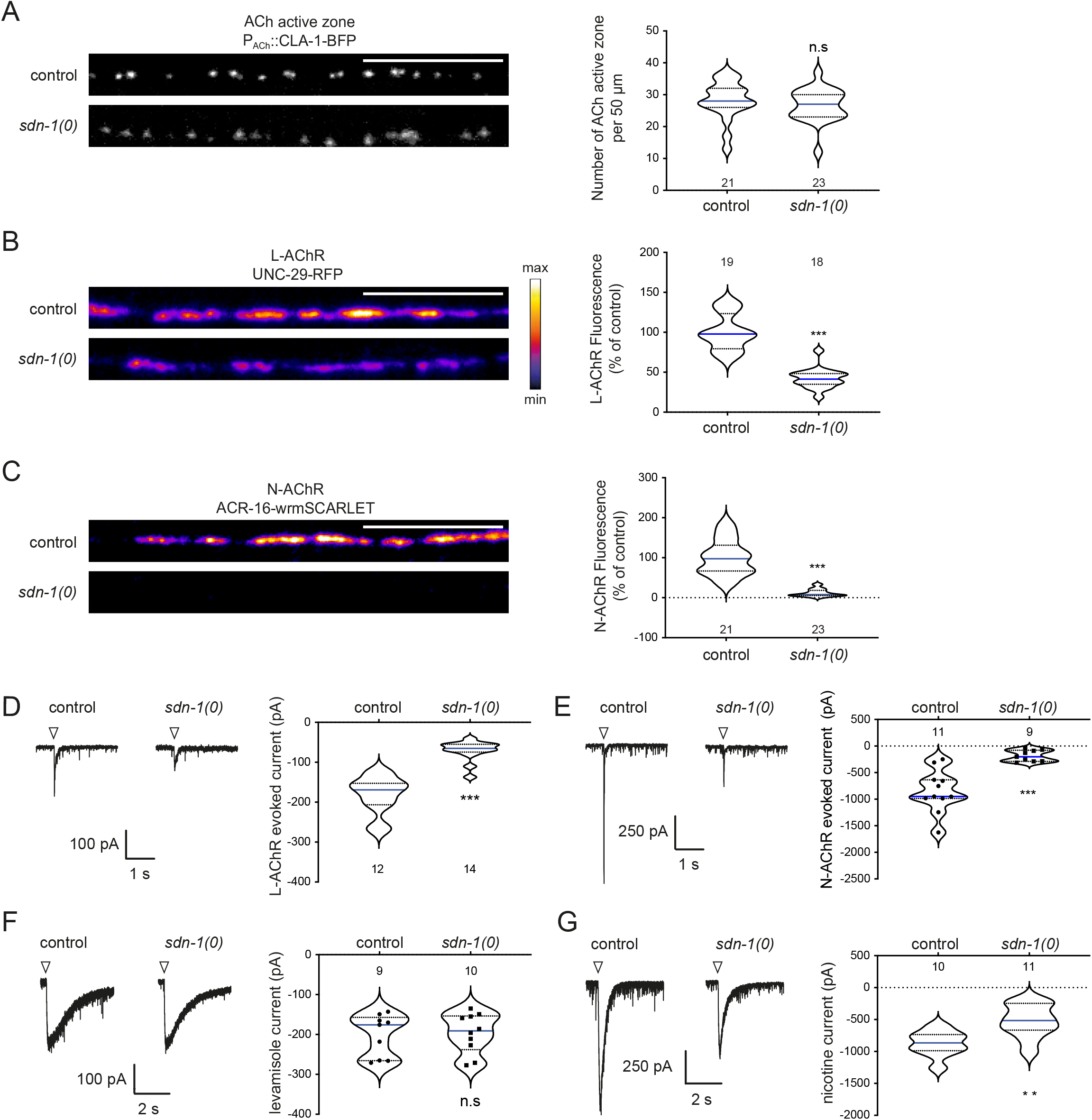
SDN-1 controls AChR synaptic content. (A) Confocal detection of the presynaptic active zone marker CLA-1-BFP under the control of the cholinergic neuron-specific promoter *Punc-17* in control and *sdn-1(0)* mutant animals. Right: quantification of active zone numbers per 50 μm of dorsal cord in control and *sdn-1(0)* animals. (B) Confocal detection and quantification of RFP-labeled L-AChR (UNC-29-RFP) fluorescence in control and *sdn-1(0)* animals. The fire image ruler represents the min to max value of intensities for each image. (C) Confocal detection and quantification of RFP-labeled N-AChR (ACR-16-wrmScarlet) fluorescence expressed from an *acr-16-wrmScarlet* knock-in allele in control and *sdn-1(0)* animals. (D-E) Representative traces of isolated L-AChR (D) or total AChR (E) peak currents evoked by 10 ms light stimulation recorded from muscle cells of control and *sdn-1(0)* animals expressing ChR2 in cholinergic motoneurons. Arrowheads mark the light stimulation onset. Isolation of L-AChR current was obtained by adding dihydro-beta-erythroidine (DHβE, 0.01 mM) in the bath solution. Quantification was done for isolated L-AChR and N-AChR currents as detailed in materials and methods. (F-G) Representative traces and quantification of peak currents evoked by 0.1 mM levamisole (F) or nicotine (G) on muscle cells from control and *sdn-1(0)* mutant animals. Arrowheads mark the 100 ms application onsets. Individual values are shown as dots in (E and F). Scale bars = 10 μm for (A, B and C). For the quantification of fluorescence levels, data were normalized to the mean value of the control group. Violin plots as in figure 1G. Mann-Whitney tests, ***p<0.001, **p<0.01, n.s: not significant. See also Fig. S2.

To confirm the synaptic localization defects of AChRs in *sdn-1(0)* mutants, we performed electrophysiological analysis. We first evoked synaptic release of acetylcholine by optogenetic stimulation of cholinergic motoneurons. The L-AChR-dependent response recorded in the body-wall muscles of *sdn-1(0)* mutants was diminished by 60 % as compared to the wild type (Fig. 2D), while the N-AChR-dependent response was reduced by 80 % (Fig. 2E). Such decrease could be due to a redistribution of receptors outside of the synapses or to a decrease of functional receptors at the plasma membrane. To test these hypotheses, we pressure-applied selective agonists of L- or N-AChRs. The response to levamisole was similar in *sdn-1(0)* and wild-type animals (Fig. 2F), indicating that in the absence of syndecan, L-AChRs are less efficiently clustered at synapses but remain present at the plasma membrane. By contrast, the response to nicotine was decreased by 40 % in *sdn-1(0)* mutants (Fig. 2G). This decrease was much smaller than the almost complete disappearance of N-AChR-dependent synaptic responses. Therefore, it suggests that syndecan has two functions. First, it is necessary for clustering N-AChRs at synapses. Second, it would stabilize N-AChRs at the plasma membrane. A simple-model would predict that N-AChRs are less stable and get endocytosed or degraded when they are not concentrated at the synapse.

Altogether, these data show that syndecan is an essential component required for proper synaptic localization of α7-like N-AChRs and, to a lesser extent, of heteromeric L-AChRs.

### Synaptic syndecan is mainly expressed by postsynaptic muscle cells

NMJs are established between neurons and muscles and are in close contact with epidermal cells. We therefore wondered which tissue contributed to the expression of syndecan at synapses. Previous studies using traditional transgenic transcriptional reporters showed that *sdn-1* was expressed in epidermis and neurons, but not in body-wall muscle cells (Minniti et al., 2004; Rhiner et al., 2005). To further investigate the *sdn-1* expression pattern, we engineered a chromosomal bi-cistronic reporter by inserting an SL2-wrmScarlet cassette in the *sdn-1* locus after the end of its coding sequence (Fig. 1A). In agreement with previous results, red fluorescence was detected in most tissues including epidermis and motoneurons, but wrmScarlet was also readily detected in body-wall muscles (Fig. 3A).

**Figure 3.**
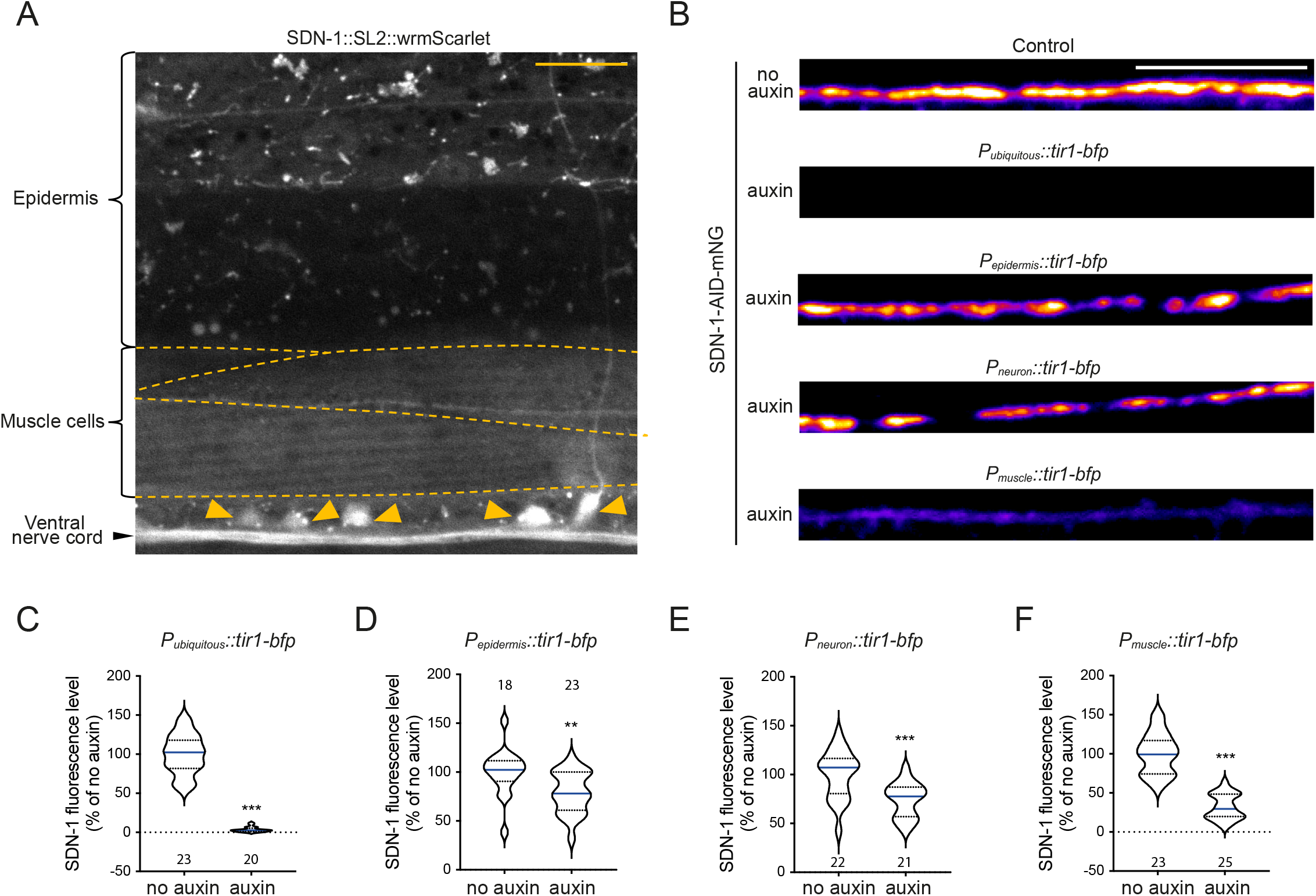
Synaptic syndecan is mainly provided by postsynaptic muscle cells. (A) Confocal detection of the transcriptional reporter SDN-1-SL2-wrmScarlet from a knock-in allele showing *sdn-1* expression in epidermis, muscle cells and nerve cord. Arrowheads indicate cell bodies of motor neuron. Dotted lines delineate borders of body-wall muscle cells. (B) Confocal detection of SDN-1-AID-mNG from dorsal nerve cord in animals grown on regular NGM plates (control) or on plates containing auxin to induce SDN-1 depletion in all tissues (animals expressing *Peft-3::tir-1-bfp*), in epidermis (animals expressing *Pdpy-7::tir-1-bfp*), in neurons (animals expressing *Prab-3::tir-1-bfp*), or in body-wall muscles (animals expressing *Pmyo-3::tir-1-bfp*). (C-F) Quantification of SDN-1-AID-mNG fluorescence levels when SDN-1 was depleted in all tissues (C), body-wall muscles (D), neurons (E) or epidermis (F). Fluorescence levels were normalized to the mean of control. Scale bars = 10 μm. Violin plots as in figure 1G. Mann-Whitney tests, ***p<0.001, **p<0.01. See also Fig. S3.

To identify the source of synaptic syndecan, we performed tissue-specific degradation of SDN-1. Using CRISPR/Cas9, we fused an Auxin-Inducible-Degron (Zhang et al., 2015) linked to mNG at the SDN-1 C-terminus (Fig. 1A) and we expressed the TIR-1 E3-ligase under various tissue-specific promoters (Fig. S3). SDN-1-AID-mNG properly localized at synapses (Fig. 3B), similarly to mNG-SDN-1 (Fig. 1C). Ubiquitous expression of TIR-1 caused a complete disappearance of fluorescence in the worm, including at the nerve cords (Fig. 3B, C). Specific degradation of SDN-1-AID-mNG either in the epidermis (Fig. 3B, D) or in neurons (Fig. 3B, E) roughly caused a 25 % reduction of the synaptic fluorescence, while degradation of SDN-1-AID-mNG in body-wall muscles caused a 70 % decrease of the synaptic fluorescence (Fig. 3B, F). Therefore, these data indicate that SDN-1 is expressed by pre-, post- and peri-synaptic cells, yet the vast majority of SDN-1 present at synaptic sites is contributed by postsynaptic muscle cells.

### Ce-Punctin/MADD-4 localizes syndecan at synapses

AChR localization at cholinergic NMJs is controlled by the anterograde synaptic organizer Ce-Punctin/MADD-4 (Pinan-Lucarré et al., 2014). *madd-4* encodes *madd-4L* (long) and *madd-4B* (short) isoforms (Fig. 4A). The long isoforms MADD-4-L are present at cholinergic NMJs and in their absence, both L- and N-AChR clusters are redistributed and no longer retained at synapses. The short isoform MADD-4B is present at both GABAergic NMJs, where it promotes GABA_A_R clustering, and cholinergic NMJs, where it prevents MADD-4L from recruiting GABA_A_Rs (Pinan-Lucarré et al., 2014). To test if MADD-4 could also regulate the localization of SDN-1, we introduced the *mNG-sdn-1* knock-in allele in various *madd-4* mutant backgrounds. mNG-SDN-1 was almost undetectable at the nerve cords of *madd-4(0)* null mutants (Fig. 4B). We then wondered if both *madd-4* isoforms were required for SDN-1 localization. *madd-4L* mutation had no impact on the localization of SDN-1, measured by the correlation with a marker of presynaptic GABA boutons, nor on its expression level (Fig. 4C-E). Genetic removal of MADD-4B changed the localization of mNG-SDN-1 without affecting its overall content at the nerve cord. Based on the labelling of GABAergic terminals, mNG-SDN-1 was no longer detectable at GABAergic synapses but relocalized at cholinergic junctions, where MADD-4L was still present (Fig. 4C-E). Hence, these results show that SDN-1 localization is controlled by the neurally-secreted extracellular matrix protein MADD-4, and that this localization can be instructed by any of the two MADD-4 isoforms.

**Figure 4.**
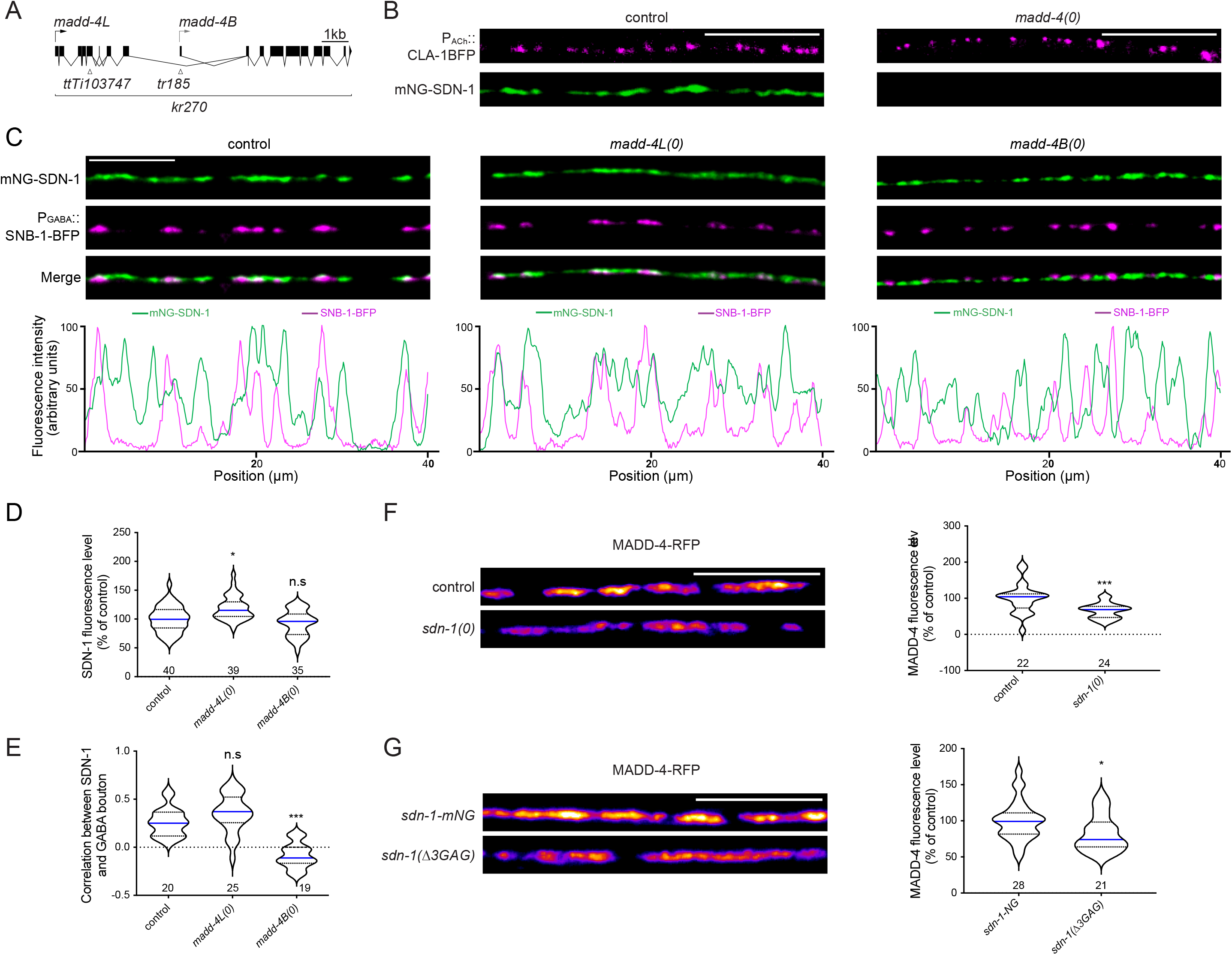
Ce-Punctin/MADD-4 localizes SDN-1 at synapses. (A) Structure of the *madd-4* locus. Boxes represent exons. Arrows indicate the beginning of the open reading frame of *madd-4L* (black) and *madd-4B* (grey). Localization of mutations are indicated: *madd-4(ttTi103747)* specifically disrupts the long isoform and is referred to as *madd-4L(0)*; *madd-4(tr185)* specifically disrupts the short isoform and is referred to as *madd-4B(0)*; *madd-4(kr270)* is a full deletion of the locus, disrupting all isoforms, and is referred to as *madd-4(0)*. (B) Confocal detection of mNG-SDN-1 and the presynaptic active zone marker CLA-1-BFP under the control of the cholinergic neuron-specific promoter *Punc-17* in control and *madd-4(0)* null animals. mNG-SDN-1 was barely detectable in *madd-4(0)* mutants. (C) Confocal detection of mNG-SDN-1 and the presynaptic marker SNB-1-BFP under the control of the GABAergic neuron-specific promoter *Punc-47* in control animals and in *madd-4L(0)* and *madd-4B(0)* mutants. The fluorescence profiles indicate mNG-SDN-1 and SNB-1-BFP fluorescence intensities along the nerve cord from the pictures above. (D) Quantification of mNG-SDN-1 fluorescence levels in control and *madd-4L(0)* and *madd-4B(0)* mutants. mNG-SDN-1 fluorescence intensities were normalized to the mean of control group. One-way ANOVA followed by Tukey’s multiple comparison tests. *p<0.05, n.s: not significant. (E) Pearson’s correlation coefficient between mNG-SDN-1 and GABA presynaptic SNB-1-BFP in control and *madd-4L(0)* and *madd-4B(0)* mutants. Kruskal-Wallis test followed by Dunn’s post-hoc test. ***p<0.001, n.s: not significant. (F-G) Confocal detection and quantification of RFP-labeled MADD-4 fluorescence in *sdn-1(0)* mutants compared to control (F) or in *sdn-1(Δ3GAG)-mNG* animals compared to control (G). Data were normalized to the mean value of the control group. Mann-Whitney tests ***p<0.001, *p<0.05. Scale bars = 10 μm. Violin plot as in figure 1G. See also Fig. S4.

We then asked whether the negatively charged GAG chains of SDN-1 might be required for its localization, because MADD-4 is predicted to contain a highly positively-charged immunoglobulin domain critical for its function (Maro et al., 2015; Seetharaman et al., 2011). SDN-1 contains two putative heparan sulfate attachment sites in its N-terminal region, and a third putative site closer to the transmembrane region (Minniti et al., 2004). We mutated these sites by CRISPR/Cas9 in a C-terminally tagged version of SDN-1, in order to avoid any potential interaction of the fluorescent protein with the extracellular matrix (Fig. S4A). Immunoprecipitation followed by western blot analysis first indicated that SDN-1 likely form SDS-resistant dimers, as previously described for mammalian syndecans (Fig. S4B) (Asundi and Carey, 1995; Choi et al., 2005). Because we did not detect any additional weight difference between the double and triple mutants, the third putative GAG attachment site is most probably not modified (Fig. S4B). Consistently, when SDN-1-mNG precipitates were treated with heparinase (I and III) the apparent molecular weight was similar as SDN-1(Δ2GAG)-mNG (Fig. S4B). Noteworthy, SDN-1(Δ3GAG)-mNG still localized at synapses, but the overall fluorescence level was decreased by 30 % when removing one chain, and by about 50 % when mutating the remaining sites as compared to the wild-type SDN-1-mNG (Fig. S4C). This suggests that GAG chains stabilize SDN-1 at the synapse, maybe through direct interaction with MADD-4, yet it is not possible to completely rule out that the point mutations introduced in the extracellular domain slightly impair the expression or the trafficking of the protein. In any case, these results demonstrate that the SDN-1 core protein carries sufficient information for MADD-4 dependent synaptic localization.

Since MADD-4 functionally interacts with SDN-1, we asked whether SDN-1 would be able, in turn, to regulate MADD-4. In *sdn-1(0)* mutants, the synaptic content of MADD-4 was reduced by 30 % as compared to the wild type, based on the expression of an RFP C-terminal fusion that labels both MADD-4 isoforms (Fig. 4F). In *sdn-1(Δ3GAG)-mNG*, MADD-4-RFP was also decreased, albeit to a lesser extent (17%) (Fig. 4G). The interpretation of these results is complicated because removing the GAG chains causes a partial loss of SDN-1, yet they suggest that the GAG chains carried by SDN-1 might partly stabilize MADD-4 at the synapse.

### SDN-1 recruits LIN-2/CASK at cholinergic synapses to cluster α7-like N-AChRs

SDN-1 contains in its C-terminus an evolutionarily conserved PDZ domain binding site, which was demonstrated to interact with the scaffolding protein CASK in the mammalian nervous system (Ethell and Yamaguchi, 1999; Hsueh et al., 1998). To determine whether the clustering of α7-like N-AChRs by SDN-1 is dependent on a SDN-1/LIN-2CASK interaction, we used CRISPR/Cas9 to delete the last 4 residues of mNG-SDN-1. The resulting protein, mNG-SDN-1(ΔEFYA), properly localized at synapses, although synaptic content was slightly decreased (Fig. S5A). However, the deletion of the PDZ-binding site caused an almost complete disappearance of N-AChRs from the nerve cord, while L-AChRs remained expressed and clustered at synapses indistinguishably from the wild type (Fig. 5A, B). We previously demonstrated that LIN-2, the ortholog of CASK that contains a PDZ domain, was present at GABAergic NMJs where it promotes the recruitment of GABA_A_Rs at GABAergic synapses (Zhou et al., 2020). Over the course of that study, we also showed that LIN-2/CASK was additionally present at cholinergic synapses. We therefore hypothesized that LIN-2 might be involved in N-AChR clustering by interacting with the SDN-1 PDZ binding site. Analysis of *acr-16-wrmScarlet* in *lin-2(0)* null mutants revealed a 95% reduction of N-AChR content at the nerve cord (Fig. 5C). This defect could be rescued by muscle specific expression of either the long isoform LIN-2A or the short isoform LIN-2B that lacks the N-terminal CaM-kinase domain but retains the PDZ, SH3 and the MAGUK domains (Fig. 5C).

**Figure 5.**
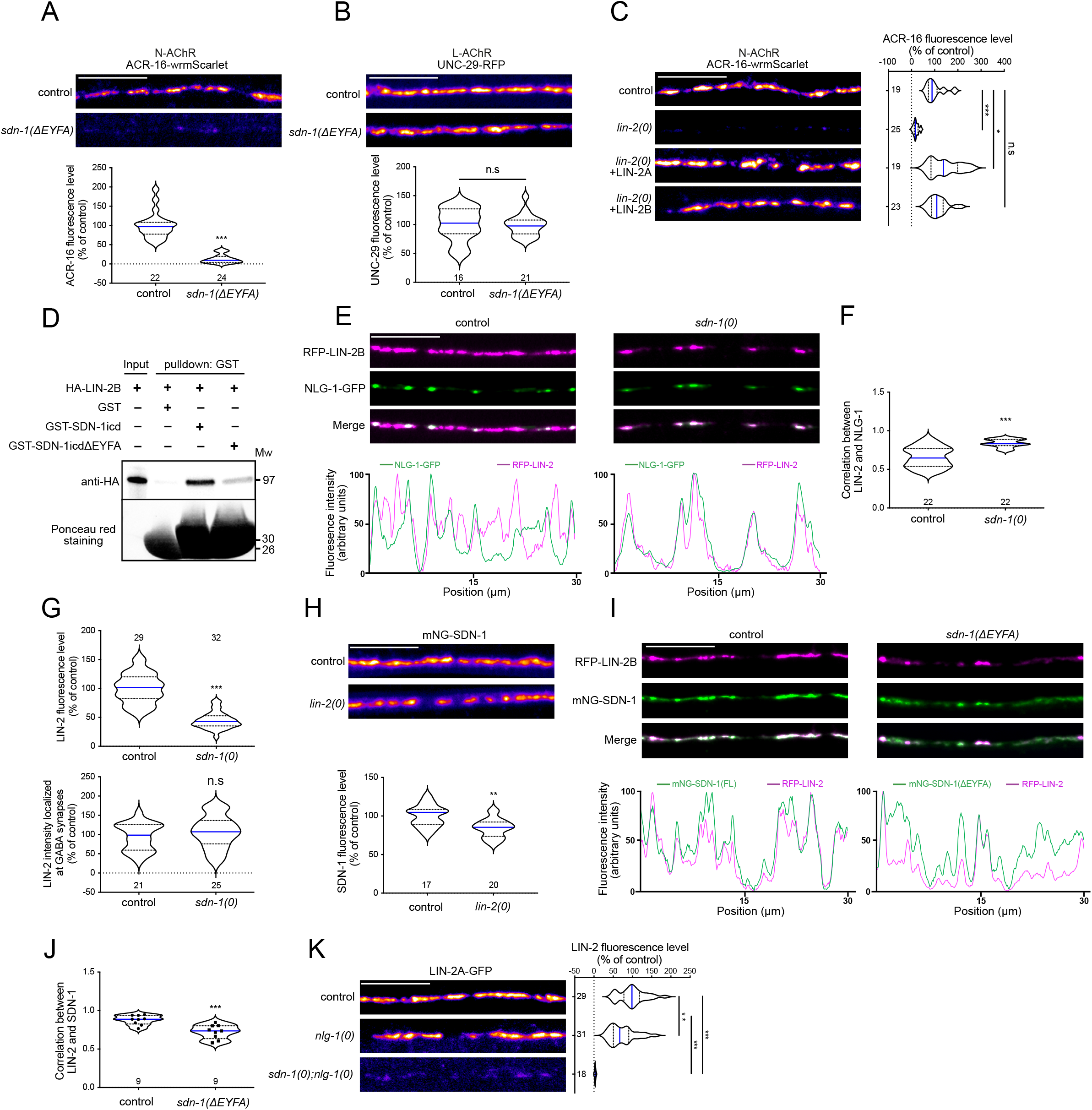
SDN-1 recruits LIN-2/CASK at cholinergic synapses to cluster α7-like N-AChRs. (A-B) Confocal detection and quantification of ACR-16-wrmScarlet (A) or UNC-29-RFP (B) fluorescence levels in control and *sdn-1(ΔEYFA)* animals that carry a mutation deleting the SDN-1 C-ter PDZ binding motif. (C) Confocal detection and quantification of ACR-16-wrmScarlet in control, *lin-2(0)*, and *lin-2(0)* animals expressing GFP-LIN-2A or GFP-LIN-2B under the control of the muscle-specific *Pmyo-3* promoter. One-way ANOVA followed by Tukey’s multiple comparison tests. ***p<0.001, *p<0.05, n.s: not significant. (D) GST pull-down analysis of LIN-2B interaction with the intracellular part of SDN-1, full size (SDN-1_ICD_) or lacking the PDZ binding motif (SDN-1_ICD_ ΔEFYA). The samples were analyzed by immunoblotting using an anti-HA antibody. The same membrane was stained with Ponceau S red to show GST expression. Molecular weights (Mw) are shown on the right. (E) Confocal detection of RFP-LIN-2B and NLG-1-GFP in control and *sdn-1(0)* animals expressing *rfp-lin-2b* and *nlg-1-gfp* under the control of the muscle-specific promoter *Pmyo-3*. The fluorescence profiles indicate RFP-LIN-2B and NLG-1-GFP fluorescence intensities along the nerve cord from the pictures above. (F) Pearson’s correlation coefficient between RFP-LIN-2B and NLG-1-GFP fluorescence. (G) Fluorescence levels of RFP-LIN-2B at the dorsal nerve cord (upper panel) or only at GABA synapses (bottom panel) in control or *sdn-1(0)* animals. Total fluorescence was measured as for other figures. To measure RFP-LIN-2B fluorescence specifically localized at GABA synapses, NLG-1-GFP marker was used to define GABA synaptic regions from processed images and RFP-LIN-2B fluorescence level was quantified in these defined regions (see material and methods for details). (H) Confocal detection and quantification of mNG-SDN-1 in control and *lin-2(0)* animals. (I) Confocal detection of RFP-LIN-2B and mNG-SDN-1 full length (control) or lacking the PDZ binding motif *(sdn-1(ΔEYFA))* in animals expressing *rfp-lin-2b* under the control of the muscle-specific promoter *Pmyo-3*. The fluorescence profiles indicate RFP-LIN-2B and mNG-SDN-1 fluorescence intensities along the nerve cord from the pictures above. (J) Pearson’s correlation coefficient between RFP-LIN-2B and mNG-SDN-1 full length or lacking the PDZ binding motif EYFA. Individual values are shown as dots. (K) Confocal detection and quantification of LIN-2A-GFP in control, *nlg-1(0)* and *sdn-1(0); nlg-1(0)* double mutants. One-way ANOVA followed by Tukey’s multiple comparison tests. ***p<0.001, **p<0.01. Scale bars = 10 μm. For the quantification of fluorescence levels, data were normalized to the mean value of the control group. Violin plot as in Figure 1G. Mann-Whitney tests for A, B, F, G, H, J: ***p<0.001, **p<0.01, *p<0.05, n.s: not significant. See also Fig. S5.

We then tested if LIN-2/CASK and SDN-1 were able to physically interact. Using *in vitro* translated and purified recombinant proteins, we showed that the intracellular domain of SDN-1 could efficiently pull-down LIN-2B. This interaction was strongly reduced after deletion of the SDN-1 PDZ-binding site (Fig. 5D). The relevance of this interaction was further tested *in vivo*. Analysis of LIN-2 expression in *sdn-1(0)* null mutants revealed a 55 % decrease of LIN-2 at the nerve cord (Fig. 5E, G). Strikingly, the remaining LIN-2 was exclusively detected at GABAergic synapses, as shown by its perfect colocalization with neuroligin/NLG-1 that is only present at GABA synapses (Maro et al., 2015; Tu et al., 2015) (Fig. 5E-G). These results suggest that SDN-1 is the limiting factor that controls the localization of LIN-2 at cholinergic synapses. By contrast, SDN-1 was only decreased by 16 % in *lin-2(0)* (Fig. 5H). Next, we analyzed the distribution of LIN-2 in *mNG-sdn-1(ΔEFYA)* mutants. LIN-2 no longer completely colocalized with truncated SDN-1 (Fig. 5I, J) and was mostly present at GABAergic NMJs (Fig. S5B-D). Because we showed that neuroligin is necessary for LIN-2 localization at GABAergic synapses, this predicts that removing both syndecan and neuroligin would cause a complete loss of LIN-2 at NMJs. Accordingly, LIN-2-GFP was almost undetectable at the nerve cord of *nlg-1(0); sdn-1(0)* double mutants (Fig. 5K).

Altogether, these data indicate that LIN-2/CASK is positioned in the postsynaptic domains of excitatory and inhibitory synapses by two distinct transmembrane proteins, namely syndecan SDN-1 and neuroligin NLG-1, respectively, that each provide binding sites for LIN-2, which in turn participates to the recruitment of AChR and GABA_A_R.

### FRM-3/FARP bridges α7-like N-AChRs with LIN-2/CASK and SDN-1

In a previous study, we showed that FRM-3, a FERM protein orthologous to mammalian FARP1/2, interacts with LIN-2 at GABAergic synapses to promote the recruitment of GABA receptors by neuroligin (Zhou et al., 2020). Because FRM-3 was also detected at cholinergic NMJs, we tested if FRM-3 might also be involved in the clustering of N-AChR. Strikingly, ACR-16-wrmScarlet was almost undetectable in a *frm-3(0)* null mutant (Fig. 6A). This phenotype could be rescued by muscle-specific expression of either full length FRM-3A or by its FERM-FA (FERM adjacent) domain. Interestingly, the FERM domain alone was not able to rescue N-AChR clustering defects, as also observed for GABA receptors at inhibitory NMJs (Fig. 6A) (Zhou et al., 2020).

**Figure 6.**
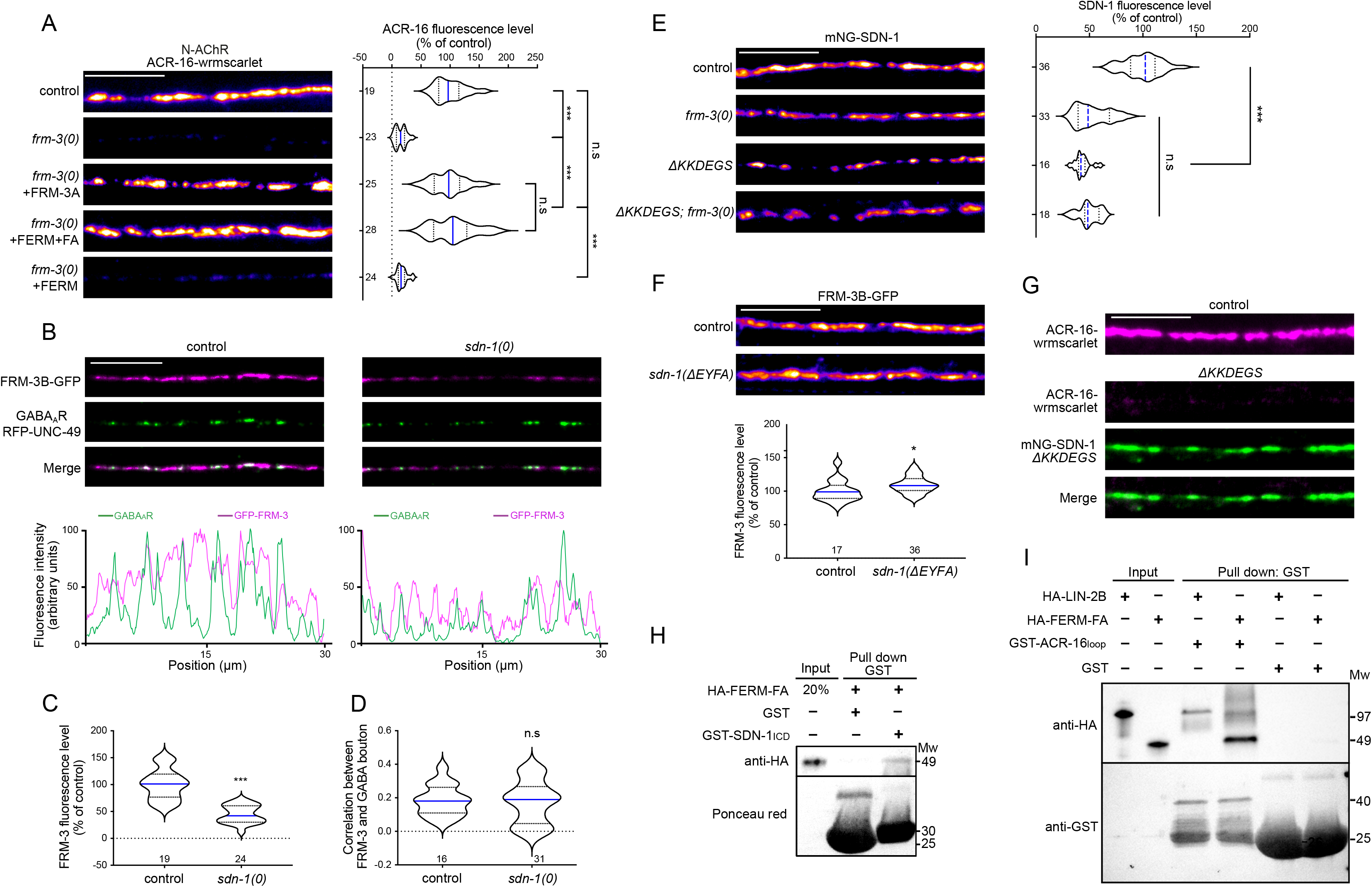
FRM-3/FARP bridges α7-like N-AChRs with LIN-2/CASK and SDN-1. (A) Confocal detection and quantification of ACR-16-wrmScarlet in control, *frm-3(0)* and *frm-3(0)* animals expressing full length FRM-3A-GFP, FERM-FA-GFP or FERM-GFP truncations under the control of the muscle-specific *Pmyo-3* promoter. (B) Confocal detection of RFP-labeled GABA_A_R (RFP-UNC-49) and FRM-3B-GFP in control and *sdn-1(0)* animals expressing *frm-3b-gfp* under the control of the muscle-specific promoter *Pmyo-3*. The fluorescence profiles indicate FRM-3B-GFP and RFP-UNC-49 fluorescence intensities along the nerve cord from the pictures above. (C) Fluorescence level of FRM-3B-GFP in control and *sdn-1(0)*. (D) Pearson’s correlation coefficient between FRM-3B-GFP and GABA bouton. (E) Confocal detection and quantification of mNG-SDN-1 full length or lacking the KKDEGS motif (*ΔKKDEGS*) in control or *frm-3(0)* animals. (F) Confocal detection and quantification of FRM-3B-GFP in control and *sdn-1(ΔEYFA)* animals expressing *frm-3b-gfp* under the control of the muscle-specific promoter *Pmyo-3*. Fluorescence levels of FRM-3B-GFP were normalized to the mean of control. (G) Confocal detection of ACR-16-wrmScarlet in control or in mNG-SDN-1 knock-in animals lacking the KKDEGS motif. (H) GST pull-down analysis of the binding between HA tagged FERM-FA domain of FRM-3B and the intracellular domain of SDN-1 (SDN-1_ICD_) by immunoblotting using anti-HA antibody. The stain-free gel was imaged by UV to show GST expression. Predicted molecular weights (Mw) are shown on the right. (I) GST pull-down analysis of ACR-16 TM3-TM4 cytosolic loop binding with HA tagged LIN-2B or FERM-FA domain of FRM-3 by immunoblotting using anti-HA antibody. The same membrane was probed by anti-GST antibody to show GST expression. Molecular weights (Mw) are shown on the right. Scale bars = 10 μm. For the quantification of fluorescence levels, data were normalized to the mean value of the control group. Violin plot as in Figure 1G. One-way ANOVA followed by Tukey’s multiple comparison tests for A and E: ***p<0.001, n.s: not significant. Mann-Whitney tests for C, D and F: ***p<0.001, *p<0.05, n.s: not significant.

We then analyzed the expression of FRM-3-GFP in a *sdn-1(0)* mutant and observed a 55 % decrease of fluorescence at the nerve cord (Fig. 6B, C). However, in contrast to LIN-2, the FRM-3 pattern remained unchanged (Fig. 6B-D). This result is consistent with our previous analysis showing that FRM-3 level was unaffected by the loss of LIN-2 (Zhou et al., 2020). Hence, despite the loss of LIN-2 at cholinergic synapses in *sdn-1(0)* mutants, FRM-3 remained partially concentrated in postsynaptic domains of cholinergic synapses. In a reciprocal experiment, we measured a 47 % decrease of SDN-1 content at the nerve cord of *frm-3(0)* mutants (Fig. 6E), much more than in *lin-2(0)* mutants. This suggested that the interactions between FRM-3 and LIN-2 with SDN-1 were different. Indeed, in contrast with LIN-2, FRM-3 content was not reduced by deleting the PDZ-binding site of SDN-1 (Fig. 6F). Interestingly, an early study reported direct binding of the FERM domain of Ezrin and Syndecan-2 through the DEGS motif present in its C1 intracellular region (Granés et al., 2003). This sequence is conserved in SDN-1 (Fig. 1A). CRISPR/Cas9 deletion of the KKDEGS sequence caused a 50 % decrease of mNG-SDN-1 fluorescence level, similar to what was observed in *frm-3(0)* mutants (Fig. 6E). The deletion of the SDN-1 KKDEGS sequence in *frm-3(0)* animals did not cause an additional decrease of mNG-SDN-1 fluorescence, suggesting that the C1 domain of SDN-1 is the site of interaction with FRM-3. We then looked at the effect of SDN-1 KKDEGS sequence deletion on N-AChRs: ACR-16-wrmScarlet fluorescence was almost undetectable in these mutants, as observed in *frm-3(0)* mutants (Fig. 6G). Therefore, FRM-3 might recruit or stabilize SDN-1 at synaptic sites through a FERM-FA/C1 interaction, which in turn recruits ACR-16.

Our data show that FRM-3 and LIN-2 are both critical for SDN-1 dependent clustering of N-AChR at cholinergic synapses. We and others previously identified physical interactions between FRM-3 and LIN-2 (Tong et al., 2015; Zhou et al., 2020). We therefore hypothesized that they might form a complex connecting N-AChR and SDN-1. Accordingly, *in vitro* pull-down experiments demonstrated interactions of the FERM-FA domain of FRM-3 with the intracellular domain of SDN-1 and with the TM3-TM4 cytoplasmic loop of ACR-16 (Fig. 6H, I). Some interaction could also be detected between LIN-2B and ACR-16 (Fig. 6I). Altogether, these data suggest that a macromolecular scaffold assembles at cholinergic synapses to localize N-AChR.

### Ce-Punctin/MADD-4 recruits UNC-40/DCC and syndecan/SDN-1 to stabilize the α7-like N-AChR ACR-16 at the synapse

We previously showed that FRM-3 is localized at NMJs by the UNC-40/DCC transmembrane receptor through a physical interaction between its FERM-FA domain and the C-terminal P3 domain of UNC-40 (Zhou et al., 2020). We therefore asked whether UNC-40 might also be required for the synaptic localization of N-AChRs. In *unc-40(0)* mutants, ACR-16-wrmScarlet fluorescence was indeed reduced by roughly 70 %, but the remaining signal had a punctate pattern (Fig. 7A). Interestingly, just deleting the P3 domain of UNC-40 caused similar loss of ACR-16, while *unc-40(ΔP3)* mutants are much healthier and have less axonal and cell migrations defects than *unc-40(0)* animals (Zhou et al., 2020).

**Figure 7.**
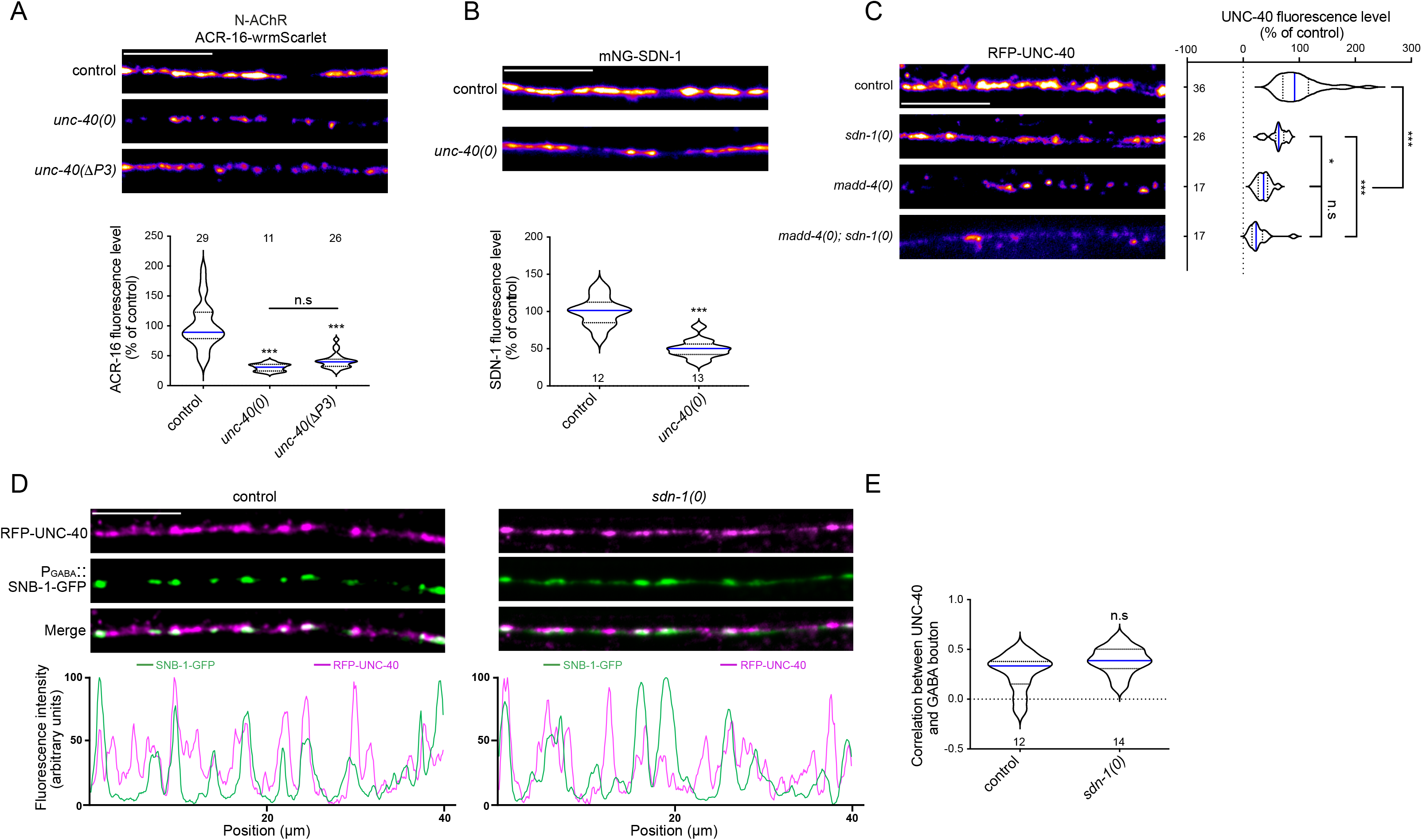
UNC-40 promotes the synaptic localization of ACR-16 and SDN-1. (A) Confocal detection and quantification of ACR-16-wrmScarlet fluorescence expressed in control, *unc-40(0)* and *unc-40(ΔP3)* mutants. Kruskal-Wallis test followed by Dunn’s post hoc test. (B) Confocal detection and quantification of mNG-SDN-1 in control and *unc-40(0)* mutant animals. Mann-Whitney test. (C) Confocal detection and quantification of RFP-UNC-40 under the control of muscle-specific *Pmyo-3* promoter in control, *sdn-1(0)*, *madd-4(0)*, and *madd-4(0); sdn-1(0)* animals. One-way ANOVA followed by Tukey’s multiple comparison tests. (D) Confocal detection of RFP-UNC-40 and the presynaptic vesicle marker SNB-1-GFP under the control of the GABAergic neuron-specific promoter *Punc-25* in control and *sdn-1(0)* animals. The fluorescence profiles indicate RFP-UNC-40 and SNB-1-GFP fluorescence intensities along the nerve cord from the pictures above. (E) Pearson’s correlation coefficient between RFP-UNC-40 and GABA boutons. Mann-Whitney test. Scale bars = 10 μm. For the quantification of fluorescence levels, data were normalized to the mean value of the control group. Violin plots as in figure 1G. ***p<0.001, *p<0.05, n.s: not significant.

Since we know that P3 interacts with FRM-3 and that the loss of FRM-3 causes a decrease of SDN-1, we predict that UNC-40 might stabilize SDN-1 via FRM-3. We therefore analyzed the SDN-1 content at the nerve cord of *unc-40(0)* mutants. In agreement with our hypothesis, we observed a 50 % loss of mNG-SDN-1 fluorescence (Fig. 7B). However, these data do not rule out a reverse interaction between SDN-1 and UNC-40, as previously reported in other systems (Bennett et al., 1997; Blanchette et al., 2015). We therefore analyzed the synaptic localization of RFP-UNC-40 in *sdn-1(0)* mutants and observed a 38 % decrease of fluorescence (Fig. 7C). Interestingly, UNC-40 remained present at both cholinergic and GABAergic NMJs (Fig. 7D, E). These results support the existence of a crosstalk between SDN-1 and UNC-40 to maintain the proper level of either protein at NMJs.

We and others previously demonstrated that UNC-40 localization at NMJs depends on the synaptic organizer MADD-4 (Tu et al., 2015). Since we demonstrated that MADD-4 localizes SDN-1 at synapses, UNC-40 localization might depend on an interaction with SDN-1. Alternatively, MADD-4 is able to bind UNC-40 in heterologous systems (Tu et al., 2015) and might therefore localize UNC-40 at synapses independently from SDN-1. To test these hypotheses, we analyzed RFP-UNC-40 expression in *madd-4(0)* mutants and observed a stronger decrease than in *sdn-1(0)* mutants (Fig. 7C), suggesting that MADD-4 has the capability to localize UNC-40 at NMJs independently from SDN-1. Consistently, the loss of UNC-40 was not aggravated in a *madd-4(0); sdn-1(0)* double mutant as compared to *madd-4(0)* single mutants (Fig. 7C).

Altogether, these results show that MADD-4 is the key-molecule that recruits SDN-1 and UNC-40 at NMJs. These three molecules then engage a network of extracellular and intracellular interactions, involving FRM-3 and LIN-2, to stabilize each other at the synapse. This, in turn, supports the recruitment of ACR-16 at cholinergic synapses.

### The intracellular domain of SDN-1 is sufficient to relocalize N-AChRs at GABAergic synapses independently from L-AChRs

At cholinergic synapses, SDN-1 is required for the localization of both L-AChRs and N-AChRs. Yet, the interaction between the intracellular C-terminus of SDN-1 and the PDZ domain of LIN-2/CASK is specifically involved in the clustering of homomeric N-AChRs, but not heteromeric L-AChRs. We therefore asked whether SDN-1 was only a permissive factor requiring additional components to specify N-AChR localization or if it was carrying sufficient information to dictate the receptor localization. To answer this question, we expressed in the muscle cells of *sdn-1(0)* mutants a GFP-tagged chimeric protein containing the extracellular and transmembrane regions of NLG-1 and the intracellular domain of SDN-1 (Fig. 8A). As predicted, this NLG-1 chimera is localized at GABA synapses (Fig. 8B) through MADD-4 interactions (Tu et al., 2015). Strikingly, as opposed to *sdn-1(0)* mutants, ACR-16-wrmScarlet were readily detected in transgenic animals but they relocalized at GABAergic NMJs together with the chimera (Fig. 8C, D). In contrast, the decreased level of L-AChRs was not rescued and the remaining receptors remained localized at cholinergic synapses (Fig. 8E-G). These results demonstrate that the intracellular domain of SDN-1 is necessary and sufficient to recruit N-AChR at synaptic sites. Furthermore, despite the fact that homomeric and heteromeric AChRs colocalize at cholinergic NMJs, their clustering relies on 2 distinct machineries (see Fig. 9 and discussion).

**Figure 8.**
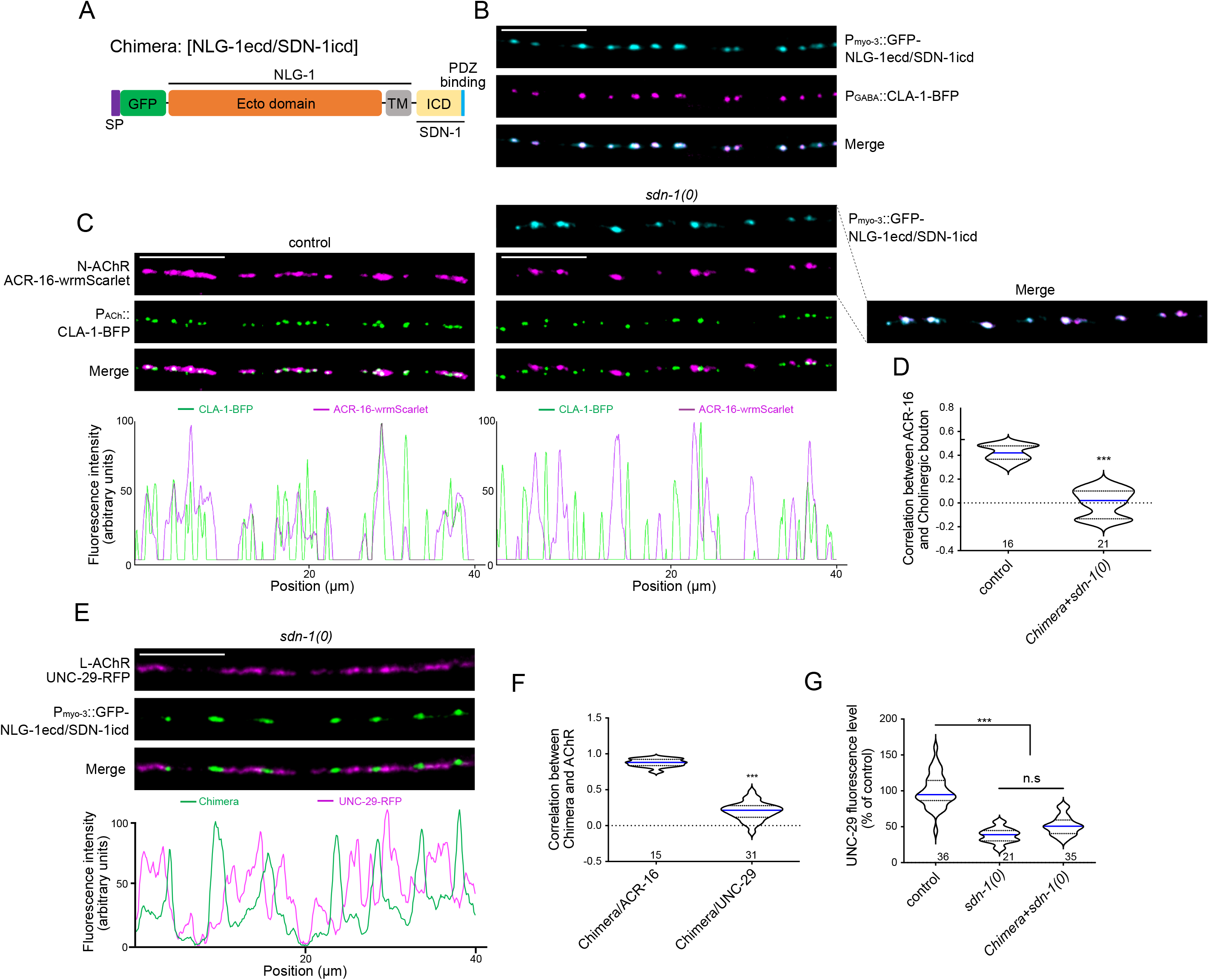
The intracellular domain of SDN-1 recruits N-AChR to GABA synapses. (A) Protein structure of the chimeric protein composed of NLG-1 ecto- and transmembrane (TM) domains and SDN-1 intracellular domain (ICD). GFP tag was inserted immediately after the signal peptide (SP). PDZ binding motif of SDN-1 is shown in blue. (B) Confocal detection of the chimeric protein GFP-NLG-1ecd/SDN-1icd expressed under the control of the muscle-specific promoter *Pmyo-3* and the presynaptic marker CLA-1-BFP under the control of the GABAergic neuron-specific promoter *Punc-47*. (C) Confocal detection of ACR-16-wrmScarlet and the presynaptic active zone marker CLA-1-BFP under the control of the cholinergic neuron-specific promoter *Punc-17* in control and *sdn-1(0)* animals. Chimeric protein NLG-1ecd/SDN-1icd was expressed under the muscle-specific promotor *Pmyo-3* in *sdn-1(0)* mutant animal. The fluorescence profiles indicate ACR-16-wrmScarlert and CLA-1-BFP fluorescence intensities along the nerve cord from the pictures above after image processing (see materials and methods for details). (D) Pearson’s correlation coefficient between N-AChRs and cholinergic boutons. Mann-Whitney test. (E) Confocal detection of RFP-labeled L-AChRs (UNC-29-RFP) and the chimeric protein GFP-NLG-1ecd/SDN-1icd expressed under the control of the muscle-specific promoter *Pmyo-3* in *sdn-1(0)* mutants. The fluorescence profiles indicate UNC-29-RFP and GFP-Chimera fluorescence intensities along the nerve cord from the pictures above. (F) Pearson’s correlation coefficient between the chimera and N-AChRs or L-AChRs in *sdn-1(0)* mutants. Mann-Whitney test. (G) Quantification of UNC-29-RFP in control, *sdn-1(0)* and *sdn-1(0)* expressing the chimera specifically in muscle. Fluorescence levels were normalized to the mean of control. Kruskal-Wallis test followed by Dunn’s post test. Scale bars = 10 μm. Violin plots as in figure 1G. ***p<0.001, n.s: not significant.

**Figure 9.**
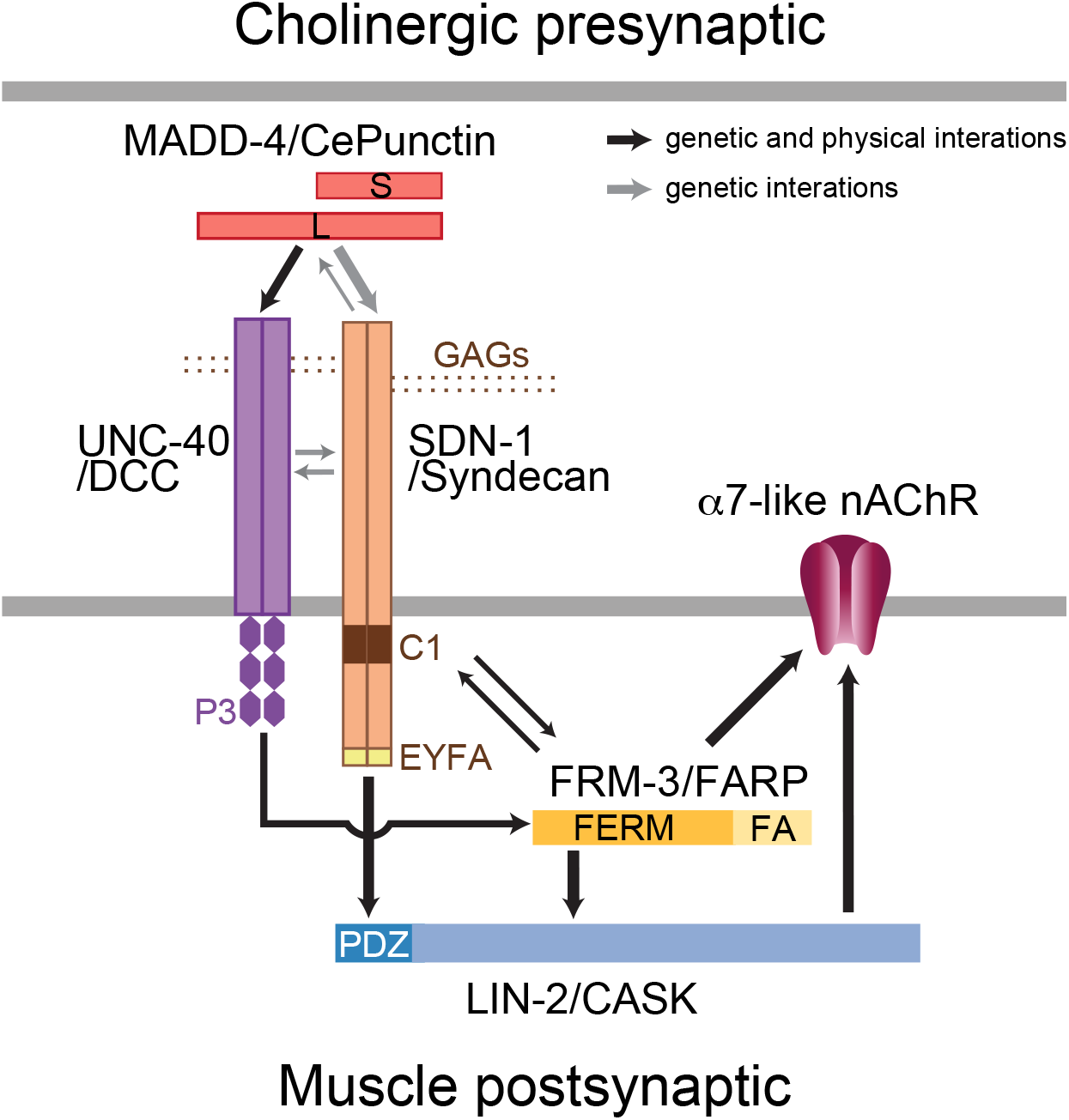
Working model for N-AChR clustering at NMJs. See discussion for details.

## DISCUSSION

Our results identified the syndecan SDN-1 as a core organizer of cholinergic and, to a lesser extent, GABAergic neuromuscular junctions in *C. elegans*. SDN-1 is absolutely required for the synaptic clustering of homomeric alpha-7-like N-AChRs and is necessary for achieving proper synaptic levels of heteromeric L-AChRs and GABA_A_ receptors. We therefore uncovered a previously unknown function of syndecan in the direct control of synaptic receptor composition. Syndecan acts by bridging extracellular matrix components with intracellular scaffolding proteins (Fig. 9). In our model, the anterograde organizer Punctin is secreted by cholinergic and GABAergic motoneurons in the synaptic cleft and triggers appropriate postsynaptic differentiation. At cholinergic synapses, Punctin has at least two parallel functions. First, it localizes the syndecan produced by muscle cells at postsynaptic sites. Second, it concentrates and likely activates the transmembrane receptor UNC-40/DCC. Syndecan, in turn, stabilizes Punctin and UNC-40 at synaptic sites. Coincident clustering of SDN-1 and UNC-40 at cholinergic NMJs triggers the intracellular recruitment of the scaffolding molecules LIN-2/CASK and FRM-3/FARP by direct interaction with the PDZ domain binding site of SDN-1 and the C-terminal P3 domain of UNC-40, respectively. The FERM-FA domain of FRM-3/FARP also engages direct interaction with SDN-1, likely with its submembrane C1 domain. The resulting CASK/FARP complex then triggers synaptic clustering of the α7-like N-AChR.

### The synaptic localization of syndecan is controlled by the ECM protein Punctin

The molecular mechanisms that control the synaptic localization of syndecan at drosophila NMJs or in rodent synapses remain elusive. At the *C. elegans* NMJ, our genetic analyses demonstrate the instructive role of Punctin for synaptic localization of SDN-1/syndecan. Strikingly, removing Punctin from GABAergic NMJs relocalizes SDN-1 exclusively at cholinergic NMJs, while in a *Punctin* null mutant, SDN-1 is almost undetectable. The most parsimonious model would predict a direct interaction between the C-terminal half of Punctin, shared by all Punctin isoforms, and SDN-1, adding to the long list of extracellular matrix syndecan interactors (see Gondelaud and Ricard-Blum, 2019). This interaction must involve the SDN-1 core protein since ΔGAG SDN-1 mutants still localize at the synapse. Accordingly, the human ADAMTS1 protease, which contain structural domains similar to Punctin, is able to cleave syndecan-4 independently of the presence of GAG chains (Rodríguez-Manzaneque et al., 2009). However, heparan sulfate (HS) chains are important to regulate the amount of SDN-1 at synapses. Using CRISPR/Cas9 in combination with biochemical analysis, we demonstrated that SDN-1 carries two HS chains. Mutation of the GAG-attachment sites causes a 50 % decrease of synaptic SDN-1. HS form long, highly negatively charged molecules able to engage versatile electrostatic interactions (Xu and Esko, 2014). Punctin is a good candidate for synaptic stabilization of SDN-1 through HS interaction because it contains an Ig-like domain with a predicted highly electropositive surface pocket, which we demonstrated to bind heparin *in vitro* (Platsaki et al, submitted). Conversely, SDN-1 tunes the synaptic amount of Punctin since we observed a 30 % decrease of Punctin in *sdn-1* null mutants. The synaptic localization mechanisms of Punctin are unknown and SDN-1 is the first identified component able to regulate the amount of Punctin. Although Punctin might interact with a yet unidentified presynaptic receptor, an alternative hypothesis is that it is locally deposited at synaptic terminals and its interaction with the ECM would prevent its diffusion and trigger the local differentiation of postsynaptic domains (Pinan-Lucarré et al., 2014). The Punctin/SDN-1 crosstalk identified at the *C. elegans* NMJ provides an interesting example of a positive feedback loop between the synaptic organizer and its effector, which likely contributes to synaptic stabilization.

### Cooperation between syndecan- and UNC-40/DCC-dependent signaling to localize α7-like N-AChRs

Syndecan impacts the second signaling output of Punctin, which involves the transmembrane receptor UNC-40/DCC. During development, Punctin promotes the growth of muscle arms toward nerve cords by positioning and activating UNC-40-dependent remodeling of the actin cytoskeleton (Alexander et al., 2009). In the adult, Punctin binds, recruits and likely activates UNC-40/DCC at the postsynaptic sites of NMJs (Tu et al., 2015). At GABAergic NMJs, UNC-40 activation sets the amount of synaptic GABA_A_Rs by promoting their recruitment to NLG-1/neuroligin (Zhou et al., 2020). Here we show that in addition to its dependence on Punctin, the synaptic content of UNC-40 is regulated by SDN-1, both at cholinergic and GABAergic synapses. In the absence of syndecan, UNC-40 remains synaptic but its content is reduced by half. DCC is known to bind heparin (Bennett et al., 1997) and numerous studies have documented the interplay between GAGs and DCC-signaling for cellular and axonal migration in various organisms (Blanchette et al., 2015, 2017; Bülow et al., 2002; Matsumoto et al., 2007; Rhiner et al., 2005). An elegant structural study demonstrated the critical role of HS for the binding of netrin to DCC by providing sulfate ions that bridge the positively charged surfaces that interact which each other on both proteins (Finci et al., 2014). Similarly, syndecan might serve as a co-receptor to strengthen the binding of Punctin on UNC-40, or stabilize UNC-40 at the plasma membrane by favoring its interaction with the ECM.

Syndecan- and DCC-dependent signaling then converge to build an intracellular scaffold containing FRM-3/FARP and LIN-2/CASK at cholinergic NMJs. This configuration parallels what we recently described at GABAergic NMJs for the adhesion molecule NLG-1/neuroligin and the transmembrane receptor UNC-40/DCC where (i) UNC-40 recruits FRM-3 through its C-terminal P3 domain, (ii) FRM-3 recruits LIN-2, (iii) LIN-2 binds the C-terminal tail of neuroligin through its PDZ domain, (iv) LIN-2 interacts with the intracellular loop of the GABA_A_R UNC-49 and stabilizes the receptors at the synapse by bridging them with Punctin-dependent neuroligin clusters (Zhou et al., 2020). At a first glance, syndecan replaces neuroligin at cholinergic synapses to eventually achieve N-AChR clustering. We demonstrated that SDN-1 interacts with LIN-2/CASK via its C-terminal PDZ domain binding site and this interaction is necessary for the localization of LIN-2/CASK at cholinergic synapses. The PDZ-dependent interaction between syndecans and CASK is evolutionarily conserved (Hsueh et al., 1998). Interestingly, CASK is mostly found in axons during development, together with SDC3, and redistributes to somatodendritic compartments in the adult, where it is coenriched with SDC2 at various synapses (Hsueh and Sheng, 1999; Hsueh et al., 1998). Our results in *C. elegans* suggest that syndecan localization might instruct the subcellular localization of CASK in mammalian neurons. As FRM-3 stabilizes SDN-1, the formation of the UNC-40-dependent intracellular scaffold potentially provides an additional level for locally-restricted stabilization of postsynaptic specialization in addition to the positive crosstalk between Punctin and syndecan in the synaptic cleft.

### The α7-like N-AChR clustering relies on an evolutionarily-conserved intracellular scaffold

The intracellular moiety of SDN-1 syndecan is the core component that controls the synaptic clustering of the alpha-7-like N-AChR by nucleating the assembly of the LIN-2/CASK-FRM-3/FARP complex at cholinergic synapses. Punctin therefore achieves N-AChR clustering by positioning SDN-1 in register with cholinergic boutons, and SDN-1 subsequently recruits N-AChRs via the LIN-2/FRM-3 cytoplasmic complex. This mechanism is fundamentally different from the clustering mechanisms of the heteromeric L-AChRs, where a set of extracellular proteins associate with L-AChRs to form small clusters that are subsequently localized at cholinergic synapses by the long isoform of Punctin (Gally et al., 2004; Gendrel et al., 2009; Pinan-Lucarré et al., 2014; Rapti et al., 2011). However, even if SDN-1 is not strictly required for L-AChR clustering, it modulates the synaptic content of L-AChRs. This likely involves extracellular interactions because neither the disruption of the SDN-1/LIN-2 association nor the ectopic expression of the SDN-1 intracellular domain has an impact on L-AChRs. The L-AChR decrease observed in *sdn-1(0)* mutants might be a consequence of the reduced amount of Punctin present at synapses. SDN-1 might also stabilize the interactions between Punctin-long and the L-AChR clustering machinery, which have still not been solved at the molecular level.

The central and specific role of SDN-1 for N-AChR clustering at cholinergic synapses raises questions about the mechanisms underlying this specificity. Not only SDN-1 is equally present at GABAergic NMJs, but its associated proteins UNC-40/DCC, FRM-3/FARP and LIN-2/CASK are used at inhibitory NMJs to cluster GABA_A_Rs. Yet, ACR-16 is exclusively detected at cholinergic synapses. Several scenarios can be envisioned. First, NLG-1 could outcompete SDN-1 for LIN-2/CASK binding and stabilize a complex that preferentially binds GABA_A_Rs. Second, Punctin-long, which is exclusively present at cholinergic NMJs, might induce distinct oligomerization of SDN-1 at the nanoscale, which is known to impact its signaling output (Choi et al., 2005; Jang et al., 2018). Third, SDN-1 might be differentially phosphorylated (Ethell et al., 2001; Horowitz and Simons, 1998; Murakami et al., 2002; Oh et al., 1997), modified or associated with a non-identified cofactor to promote the formation of an ACR-16 binding complex. The insertion of ACR-16 in the muscle membrane was shown to be regulated by Wnt ligands possibly secreted from cholinergic neurons (Babu et al., 2011; Francis et al., 2005; Jensen et al., 2012; Tikiyani et al., 2018). A crosstalk between Wnts and syndecan has been documented in many systems (Pataki et al., 2015) and SDN-1 might serve as a co-receptor at cholinergic NMJs. Yet, this putative Wnt-syndecan crosstalk can be bypassed by overexpression of the SDN-1 tail at GABAergic synapses, which emphasizes the central role of SDN-1 as a postsynaptic organizer.

The *C. elegans* ACR-16 is strikingly close to the mammalian α7 nAChR subunit (Ballivet et al., 1996; Touroutine et al., 2005). These receptors have attracted attention because they have been implicated in cognition and memory and their dysfunction was associated to neuropsychiatric disorders (for review see (Dineley et al., 2015)). In some instances, they mediate fast synaptic transmission, as in the hippocampus (Alkondon et al., 1997; Bürli et al., 2010; Fabian-Fine et al., 2001; Gray et al., 1996; Wonnacott, 1997; Zarei et al., 1999), and in many cases, they localize presynaptically on GABAergic or glutamatergic terminals where they enhance neurotransmitter release (Alkondon et al., 1997; Bürli et al., 2010; Fabian-Fine et al., 2001; Gray et al., 1996; Liu et al., 2001). Surprisingly, the mechanisms that regulate these localizations remain mostly unknown. Miscellaneous factors including synaptic activity, neurotrophin, anti-apoptotic Bcl-2 proteins, src kinase, Wnt and Neuroregulin signaling pathways or plasma membrane lipid composition (Brusés et al., 2001; Charpantier et al., 2005; Dawe et al., 2019; Farías et al., 2007; Hancock et al., 2008; Kawai et al., 2002), have been shown to impact α7 receptor localization, but always partially. Colocalization between α7 receptor clusters and PSD-95 family members has been described, yet direct interaction was not documented (Conroy et al., 2003; Farías et al., 2007; Parker et al., 2004). Because all the components we identified in *C. elegans* are evolutionarily conserved and expressed in neurons, it would be worth testing if syndecan regulates the localization of nicotinic receptors in the mammalian brain.

## ACKNOWLEDGEMENTS

We thank the Hobert laboratory for plasmids, the *Caenorhabditis* Genetic Center (which is funded by NIH Office of Research Infrastructure Programs, P40 OD010440) for strains, Alexis Weinreb for image analysis scripts, the CIQLE Imaging facility for support and access to equipment. This work was supported by the European Research Council (ERC_Adg C.NAPSE #695295), within the framework of the LABEX CORTEX (ANR-11-LABX-0042) of Université de Lyon, within the program “Investissements d’Avenir” (ANR-11-IDEX-0007) operated by the French National Research Agency (ANR). Work in the Bülow lab is supported by a grant from the National Institute of Neurological Disorders and Stroke (R21NS111145).

## AUTHOR CONTRIBUTION

Conceptualization, J.L.B., X.Z., C.V., H.E.B.; Methodology, X.Z., C.V., M.C., O.R., M.J.; Investigation, X.Z., C.V., M.C., O.R., M.J.; Writing –Original Draft, J.L.B, Z.X., C.V., M.J.; Writing –Review & Editing, M.C., H.E.B.; Funding Acquisition, J.LB., H.E.B..

## DECLARATION OF INTERESTS

The authors declare no competing interests.

## METHODS

### Strains and genetics

All *C. elegans* strains were originally derived from the wild-type Bristol N2 strain. Worm cultivation, genetic crosses, and manipulation of *C. elegans* were carried out according to standard protocols (Brenner, 1974). All strains were maintained on nematode growth medium (NGM) agar plates with *Escherichia coli* OP50 as a food source at 20 °C. A complete list of strains used in this study can be found in Table 1.

**Table 1:** List of strains.

| Figure | Strain | Genotype |
| --- | --- | --- |
| 1, 5, 6, 7, S5 | EN437 | *kr437 [mNG-sdn-1] (X)* |
| 1, S1 | EN7838 | *kr296 [unc-49-TagRFP-T] (III) ; kr437 [mNG-sdn-1] (X)* |
| 1, S1, 5 | EN7839 | *kr208 [unc-29-TagRFP-T] (I) ; kr437 [mNG-sdn-1] (X)* |
| 1, S1 | EN7860 | *krSi144 [Punc-17::cla-1-TagBFP] (I) ; kr437 [mNG-sdn-1] (X)* |
| 1, S1 | EN7915 | *krSi143 [Punc-47::cla-1-TagBFP] (V) ; kr437 [mNG-sdn-1] (X)* |
| 2 | EN7849 | *kr440 [acr-16-wrmScarlet] (I) ; krSi144 [Punc-17::cla-1-TagBFP] (I) ; sdn-1(zh20) (X)* |
| 2 | EN7850 | *kr440 [acr-16-wrmScarlet] (I) ; krSi144 [Punc-17::cla-1-TagBFP] (I)* |
| 2, S1 | EN208 | *kr208 [unc-29-TagRFP-T] (I)* |
| 2 | EN2649 | *kr208 [unc-29-TagRFP-T] (I) ; sdn-1(zh20) (X)* |
| 2 | EN7244 | *zxIs6 [Punc-17::Chr2(H134R)-YFP+lin-15(+)] (V)* |
| 2 | EN7243 | *zxIs6 [Punc-17::Chr2(H134R)-YFP+lin-15(+)] (V) ; sdn-1(zh20) (X)* |
| S1 | EN7808 | *krSi144 [Punc-17::cla-1-TagBFP] (I)* |
| S1 | EN7809 | *krSi143 [Punc-47::cla-1-TagBFP] (V)* |
| S2 | EN3886 | *kr296 [unc-49-TagRFP-T] (III) ; sdn-1(zh20) (X) ; krls67 [Punc-47::snb-1-TagBFP]* |
| S2 | EN3796 | *kr296 [unc-49-TagRFP-T] (III) ; krls67 [Punc-47::snb-1-TagBFP]* |
| S2 | EN3513 | *kr296 [unc-49-TagRFP-T] (III) ; sdn-1(zh20) (X)* |
| S1, S2 | EN296 | *kr296 [unc-49-TagRFP-T] (III)* |
| 3 | EN387 | *kr387 [sdn-1::SL2-wrmScarlet] (X)* |
| 3 | EN7094 | *kr208 [unc-29-TagRFP-T] (I) ; krSi50 [Peft-3::TIR1-TagBFP] (IV) ; kr383 [sdn-1::AID-mNG] (X)* |
| 3 | EN7158 | *kr208 [unc-29-TagRFP-T] (I) ; krSi55 [Pmyo-3::TIR1-TagBFP] (V) ; kr383 [sdn-1::AID-mNG] (X)* |
| 3 | EN7029 | *kr208 [unc-29-TagRFP-T] (I) ; krSi36 [Prab-3::TIR1-TagBFP] (V) ; kr383 [sdn-1::AID-mNG] (X)* |
| 3 | EN7235 | *kr208 [unc-29-TagRFP-T] (I) ; krSi63 [Pdpy-7::TIR1-TagBFP] (II) ; kr383 [sdn-1::AID-mNG] (X)* |
| S3 | EN7131 | *ieSi58 [Peft-3::AID-GFP::unc-54 3'UTR + Cbr-unc-119(+)] (IV) ; krSi50 [Peft-3::TIR1-TagBFP] (IV)* |
| S3 | EN7157 | *ieSi58 [Peft-3::AID-GFP::unc-54 3'UTR + Cbr-unc-119(+)] (IV) ; krSi55 [Pmyo-3::TIR1-TagBFP] (V)* |
| S3 | EN7028 | *ieSi58 [Peft-3::AID-GFP::unc-54 3'UTR + Cbr-unc-119(+)] (IV) ; krSi36 [Prab-3::TIR1-TagBFP] (V)* |
| S3 | EN7236 | *krsi63 [Pdpy-7::TIR1-TagBFP] (II) ; ieSi58 [Peft-3::AID-GFP::unc-54 3'UTR + Cbr-unc-119(+)] (IV)* |
| 4 | EN7937 | *krSi145 [Punc-17::cla-1-TagBFP] (IV) ; kr437 [mNG-sdn-1] (X)* |
| 4 | EN7938 | *madd-4(kr270) (I) ; krSi145 [Punc-17::cla-1-TagBFP] (IV) ; kr437 [mNG-sdn-1] (X)* |
| 4 | EN7747 | *kr373 [madd-4-TagRFP-T] (I) ; kr437 [mNG-sdn-1] (X) ; krls67 [Punc-47::snb-1-TagBFP]* |
| 4 | EN7754 | *madd-4(tr185) (I) ; kr437 [mNG-sdn-1] (X) ; krSi67 [Punc-47::snb-1-TagBFP]* |
| 4 | EN7755 | *madd-4(ttTi103747) (I) ; kr437 [mNG-sdn-1] (X) ; krSi67 [Punc-47::snb-1-TagBFP]* |
| 4 | EN373 | *kr373 [madd-4-TagRFP-T] (I)* |
| 4 | EN7899 | *kr373 [madd-4-TagRFP-T] (I) ; sdn-1(zh20) (X)* |
| 4 | EN7631 | *kr373 [madd-4-TagRFP-T] (I) ; kr383 [sdn-1-AID-mNG] (X)* |
| 4 | EN7758 | *kr373 [madd-4-TagRFP-T] (I) ; kr475 [sdn-1(S71A,S86A,S214A)-AID-mNG] (X)* |
| S4 | EN383 | *kr383 [sdn-1-AID-mNG] (X)* |
| S4 | EN441 | *kr441 [sdn-1(S71A)-AID-mNG]* |
| S4 | EN388 | *kr388 [sdn-1(S71A,S86A)-AID-mNG]* |
| S4 | EN475 | *kr475 [sdn-1(S71A,S86A,S214A)-AID-mNG]* |
| S5 | EN496 | *kr496 [mNG-sdn-1(ΔEYFA)] (X)* |
| S5 | EN7858 | *krSi143 [Punc-47::cla-1-TagBFP] V ; krSi35 [Pmyo-3::lin-2-RFP]* |
| S5 | EN7859 | *krSi143 [Punc-47::cla-1-TagBFP] V ; krSi35 [Pmyo-3::lin-2-RFP] ; kr496 [mNG-sdn-1(ΔEYFA)] (X)* |
| 5, 6, 7 | EN440 | *kr440 [acr-16-wrmScarlet] (I)* |
| 5 | EN7556 | *kr440 [acr-16-wrmScarlet] (I) ; kr437 [mNG-sdn-1] (X)* |
| 5 | EN7840 | *kr440 [acr-16-wrmScarlet] (I) ; kr496 [mNG-sdn-1(ΔEYFA)] (X)* |
| 5 | EN7866 | *kr208 [unc-29-TagRFP-T] (I) ; kr496 [mNG-sdn-1(ΔEYFA)] (X)* |
| 5 | EN7729 | *kr440 [acr-16-wrmScarlet] (I) ; lin-2 (n397) (X)* |
| 5 | EN7903 | *krSi30 [Pmyo-3::lin-2a-GFP] ; kr440 [acr-16-wrmScarlet] (I) ; lin-2 (n397) (X)* |
| 5 | EN7960 | *krSi60 [Pmyo-3::lin-2b-GFP] ; kr440 [acr-16-wrmScarlet] (I) ; lin-2 (n397) (X)* |
| 5 | EN7057 | *krSi35 [Pmyo-3::lin-2a-RFP] ; krSi10 [Pmyo-3::nlg-1-GFP]* |
| 5 | EN7779 | *krSi35[Pmyo-3::lin-2a-RFP] ; krSi10 [Pmyo-3::nlg-1-GFP] ; sdn-1(zh20) (X)* |
| 5 | EN7720 | *kr437 [mNG-sdn-1] (X) ; lin-2(n397) (X)* |
| 5 | EN7858 | *krSi35 [Pmyo-3::lin-2a-RFP] ; krSi143 [Punc-47::cla-1-TagBFP] ; kr437 [mNG-sdn-1] (X)* |
| 5 | EN7859 | *krSi35 [Pmyo-3::lin-2a-RFP] ; krSi143 [Punc-47::cla-1-TagBFP] ; kr437 [mNG-sdn-1] (X) ; kr496 [mNG-sdn-1(ΔEYFA)] (X)* |
| 5 | EN3921 | *krSi30[Pmyo-3::lin-2a-GFP] ; krls67 [Punc-47::snb-1-TagBFP] ; kr296 [unc-49-TagRFP-T] (III)* |
| 5 | EN7013 | *krSi30[Pmyo-3::lin-2a-GFP] ; krls67 [Punc-47::snb-1-TagBFP] ; kr296 [unc-49-TagRFP-T] (III) ; nlg-1 (ok259) (X)* |
| 5 | EN7832 | *krSi30[Pmyo-3::lin-2a-GFP] ; krls67 [Punc-47::snb-1-TagBFP] ; kr296 [unc-49-TagRFP-T] (III) ; nlg-1 (ok259)] (X) ; sdn-1(zh20) (X)* |
| 6 | EN7489 | *kr440 [acr-16-wrmScarlet] (I) ; frm-3(gk585) (X)* |
| 6 | EN7959 | *krSi31 [Pmyo-3::GFP-frm-3a] ; kr440 [acr-16-wrmScarlet] (I) ; frm-3(gk585) (X)* |
| 6 | EN8055 | *krSi32 [Pmyo-3::GFP-frm-3a(1-412aa)] ; kr440 [acr-16-wrmScarlet] (I) ; frm-3(gk585) (X)* |
| 6 | EN7912 | *krSi39 [Pmyo-3::GFP-frm-3a(1-318aa)] ; kr440 [acr-16-wrmScarlet] (I) ; frm-3(gk585) (X)* |
| 6 | EN8048 | *kr440 [acr-16-wrmScarlet] (I) ; kr437 [mNG-sdn-1] (X) ; frm-3(gk585) (X)* |
| 6 | EN50519 | *sdn-1 [mNG-kr519(ΔKKDEGS)] (X)* |
| 6 | EN8057 | *sdn-1 [mNG-kr519(ΔKKDEGS)] (X) ; frm-3(gk585) (X)* |
| 6 | EN8082 | *kr440 [acr-16-wrmScarlet] (I) ; sdn-1 [mNG-kr519(ΔKKDEGS)] (X)* |
| 6 | EN7177 | *krSi61 [Pmyo-3::frm-3-GFP] ; kr296 [unc-49-TagRFP-T] (III)* |
| 6 | EN8046 | *krSi144 [Punc-17::cla-1-TagBFP] ; krSi61 [Pmyo-3::frm-3-GFP] ; kr499 [sdn-1(ΔEYFA)] (X)* |
| 6 | EN7208 | *krSi61 [Pmyo-3::frm-3-GFP] ; krls67 [Punc-47::snb-1-TagBFP] ; kr296 [unc-49-TagRFP-T] (III)* |
| 6 | EN7209 | *krSi61 [Pmyo-3::frm-3-GFP] ; krls67 [Punc-47::snb-1-TagBFP] ; kr296 [unc-49-TagRFP-T] (III) ; sdn-1(zh20) (X)* |
| 7 | EN7512 | *unc-40(e1430) (I) ; kr440 [acr-16-wrmScarlet] (I)* |
| 7 | EN7910 | *krSi145 [Punc-17::cla-1-TagBFP] ; kr457[unc-40(ΔP3)] (I) ; kr440 [acr-16-wrmScarlet] (I)* |
| 7 | EN7733 | *unc-40(e1430) (I) ; kr437 [mNG-sdn-1] (X)* |
| 7 | EN7833 | *krSi11 [Pmyo-3::TagRFP-T-unc-40] (II) ; juls1 [Punc-25 ::snb-1-GFP] ; sdn-1(zh20) (X)* |
| 7 | EN3333 | *krSi11 [Pmyo-3::RFP-unc-40] (II)* |
| 7 | EN3361 | *krSi11 [Pmyo-3::RFP-unc-40] (II) ; juls1 [Punc-25 ::snb-1-GFP]* |
| 7 | EN7845 | *krSi11 [Pmyo-3::TagRFP-T-unc-40] (II) ; madd-4(kr270) (I)* |
| 7 | EN7831 | *krSi11 [Pmyo-3::Tag-RFP-T-unc-40] (II) ; madd-4(kr270) (I) ; sdn-1(zh20) (X)* |
| 8 | EN8107 | *krSi157[Pmyo-3::GFP-NLG-1ecd/SDN-1/icd] ; krSi143 [Punc-47::cla-1-TagBFP] ; kr296 [unc-49-TagRFP-T] (III)* |
| 8 | EN7926 | *krSi157[Pmyo-3::GFP-NLG-1ecd/SDN-1/icd] ; krSi145 [Punc-17::cla-1-TagBFP] ; kr440 [acr-16-wrmScarlet] (I) ; sdn-1(zh20) (X)* |
| 8 | EN7851 | *krSi144 [Punc-17::cla-1-TagBFP] ; kr208 [unc-29-TagRFP-T] (I) ; sdn-1(zh20) (X)* |
| 8 | EN8040 | *krSi160[Pmyo-3::GFP-NLG-1ecd/SDN-1/icd] ; kr208 [unc-29-TagRFP-T] (I) ; sdn-1(zh20) (X)* |

### Plasmids

The constructs created in this study are described in Table 2. The full open reading frames of all constructs were verified by Sanger sequencing from GATC Company. The other plasmids used to create single-copy insertion alleles by miniMos method are described in a previous study (Zhou et al., 2020).

**Table 2:**
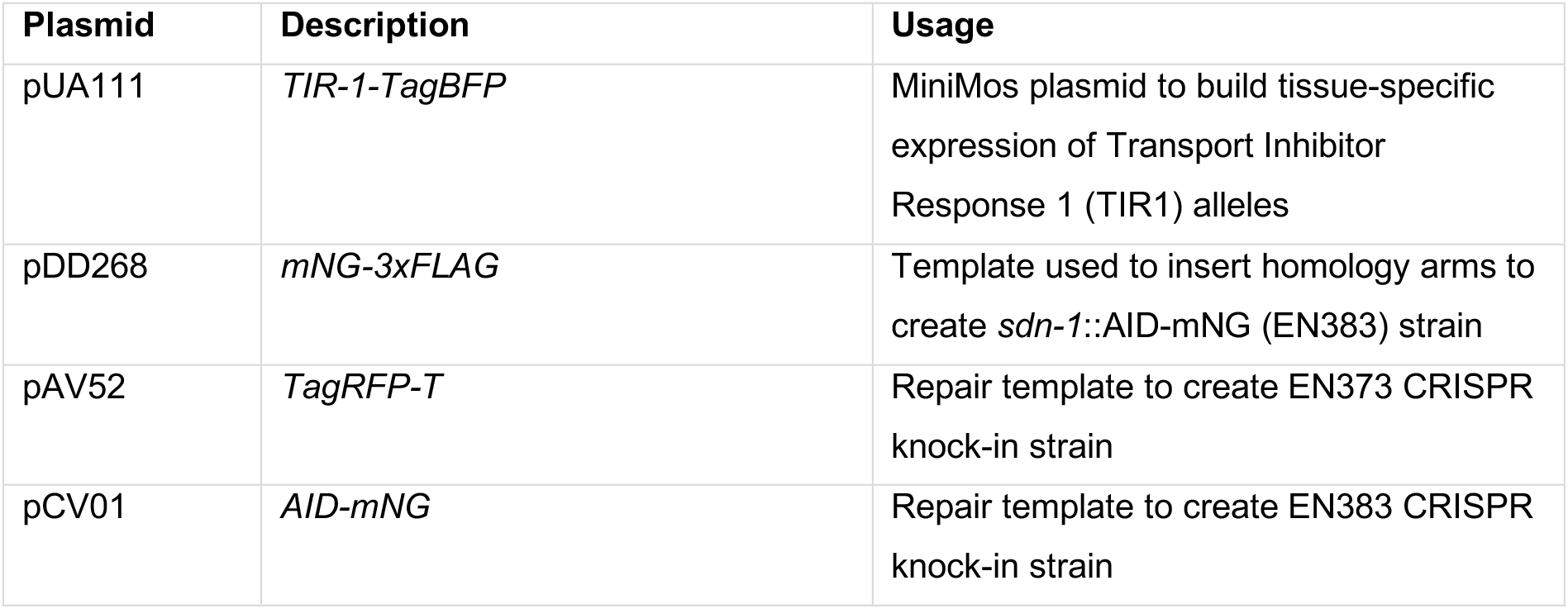
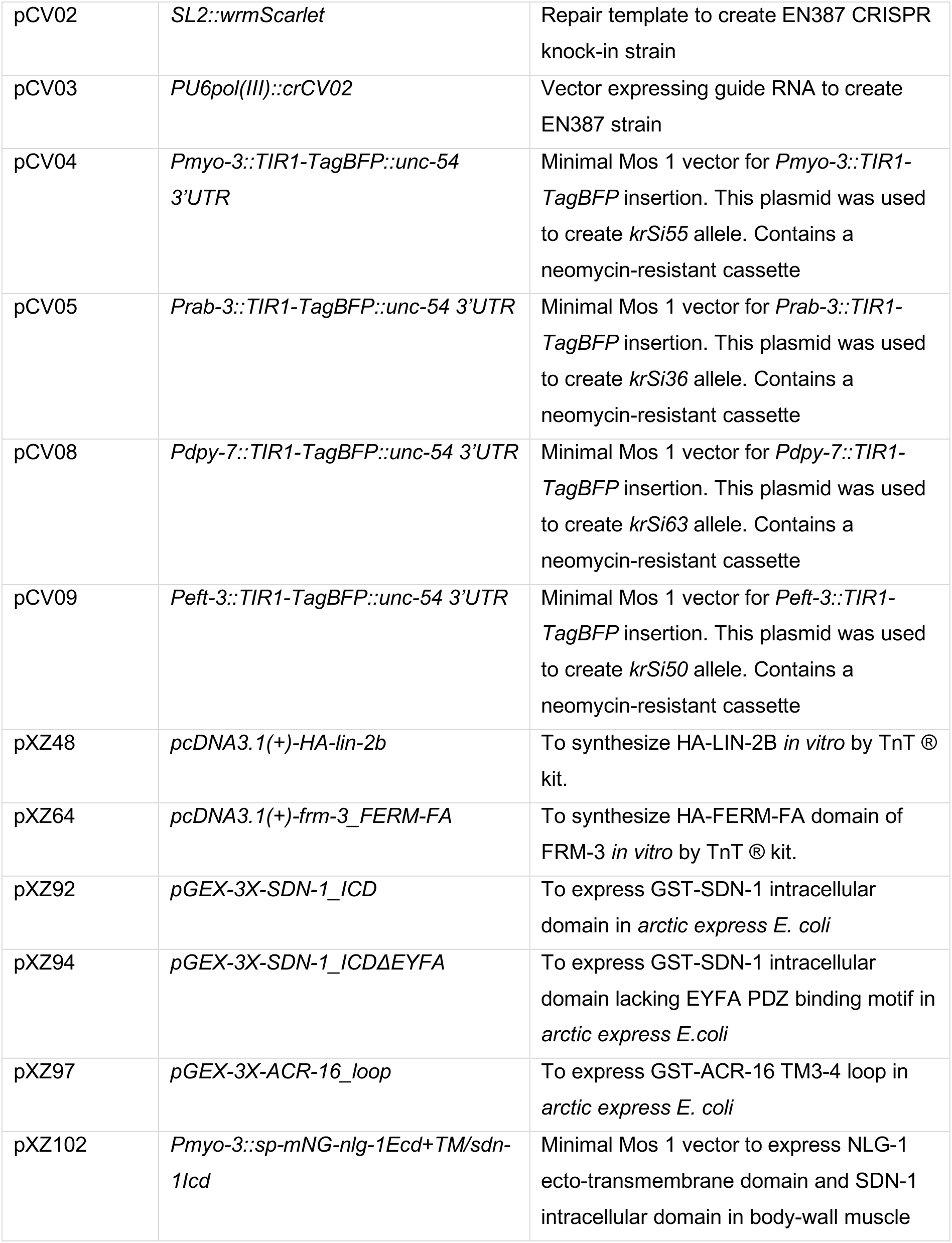
List of generated plasmids.

| Plasmid | Description | Usage |
| --- | --- | --- |
| pUA111 | *TIR-1-TagBFP* | MiniMos plasmid to build tissue-specific expression of Transport Inhibitor Response 1 (TIR1) alleles |
| pDD268 | *mNG-3xFLAG* | Template used to insert homology arms to create *sdn-1::AID-mNG* (EN383) strain |
| pAV52 | *TagRFP-T* | Repair template to create EN373 CRISPR knock-in strain |
| pCV01 | *AID-mNG* | Repair template to create EN383 CRISPR knock-in strain |
| pCV02 | *SL2::wrmScarlet* | Repair template to create EN387 CRISPR knock-in strain |
| pCV03 | *PU6pol(III)::crCV02* | Vector expressing guide RNA to create EN387 strain |
| pCV04 | *Pmyo-3::TIR1-TagBFP::unc-54 3'UTR* | Minimal Mos 1 vector for *Pmyo-3::TIR1-TagBFP* insertion. This plasmid was used to create *krSi55* allele. Contains a neomycin-resistant cassette |
| pCV05 | *Prab-3::TIR1-TagBFP::unc-54 3'UTR* | Minimal Mos 1 vector for *Prab-3::TIR1-TagBFP* insertion. This plasmid was used to create *krSi36* allele. Contains a neomycin-resistant cassette |
| pCV08 | *Pdpy-7::TIR1-TagBFP::unc-54 3'UTR* | Minimal Mos 1 vector for *Pdpy-7::TIR1-TagBFP* insertion. This plasmid was used to create *krSi63* allele. Contains a neomycin-resistant cassette |
| pCV09 | *Peft-3::TIR1-TagBFP::unc-54 3'UTR* | Minimal Mos 1 vector for *Peft-3::TIR1-TagBFP* insertion. This plasmid was used to create *krSi50* allele. Contains a neomycin-resistant cassette |
| pXZ48 | *pcDNA3.1(+)-HA-lin-2b* | To synthesize HA-LIN-2B *in vitro* by TnT ® kit. |
| pXZ64 | *pcDNA3.1(+)-frm-3_FERM-FA* | To synthesize HA-FERM-FA domain of FRM-3 *in vitro* by TnT ® kit. |
| pXZ92 | *pGEX-3X-SDN-1_ICD* | To express GST-SDN-1 intracellular domain in *arctic express E. coli* |
| pXZ94 | *pGEX-3X-SDN-1_ICDΔEYFA* | To express GST-SDN-1 intracellular domain lacking EYFA PDZ binding motif in *arctic express E.coli* |
| pXZ97 | *pGEX-3X-ACR-16_loop* | To express GST-ACR-16 TM3-4 loop in *arctic express E. coli* |
| pXZ102 | *Pmyo-3::sp-mNG-nlg-1Ecd+TM/sdn-1lcd* | Minimal Mos 1 vector to express NLG-1 ecto-transmembrane domain and SDN-1 intracellular domain in body-wall muscle |

### Generation of single-copy insertion alleles

The single-copy insertion alleles generated by miniMos method (Frøkjær-Jensen et al., 2014) are listed in Table 3. Genes encoding fluorescent tagged proteins are driven by *myo-3* (body-wall muscle), *unc-47* (GABAergic motoneuron), *unc-17* (cholinergic motoneuron), *rab-3* (pan-neuronal), *dpy-7* (epidermis), or *eft-3* (universal) specific promoters. N2 animals were injected with 15 ng/μL of plasmid of interest containing the promoter and the open reading frame, 50 ng/μL pCFJ601 (Mos1 transposase), 10 ng/μL pMA122 (negative selective marker *Phsp16.2::peel-1*), 2.5 ng/μL pCFJ90 (*Pmyo-2::mCherry*). Neomycin (G418) was added to plates 24 hours after injection at 1.5 μg/μL final concentration. Candidate plates were heat-shocked for 2 hours at 34°C to remove extrachromosomal arrays and then homozygosed.

**Table 3:**
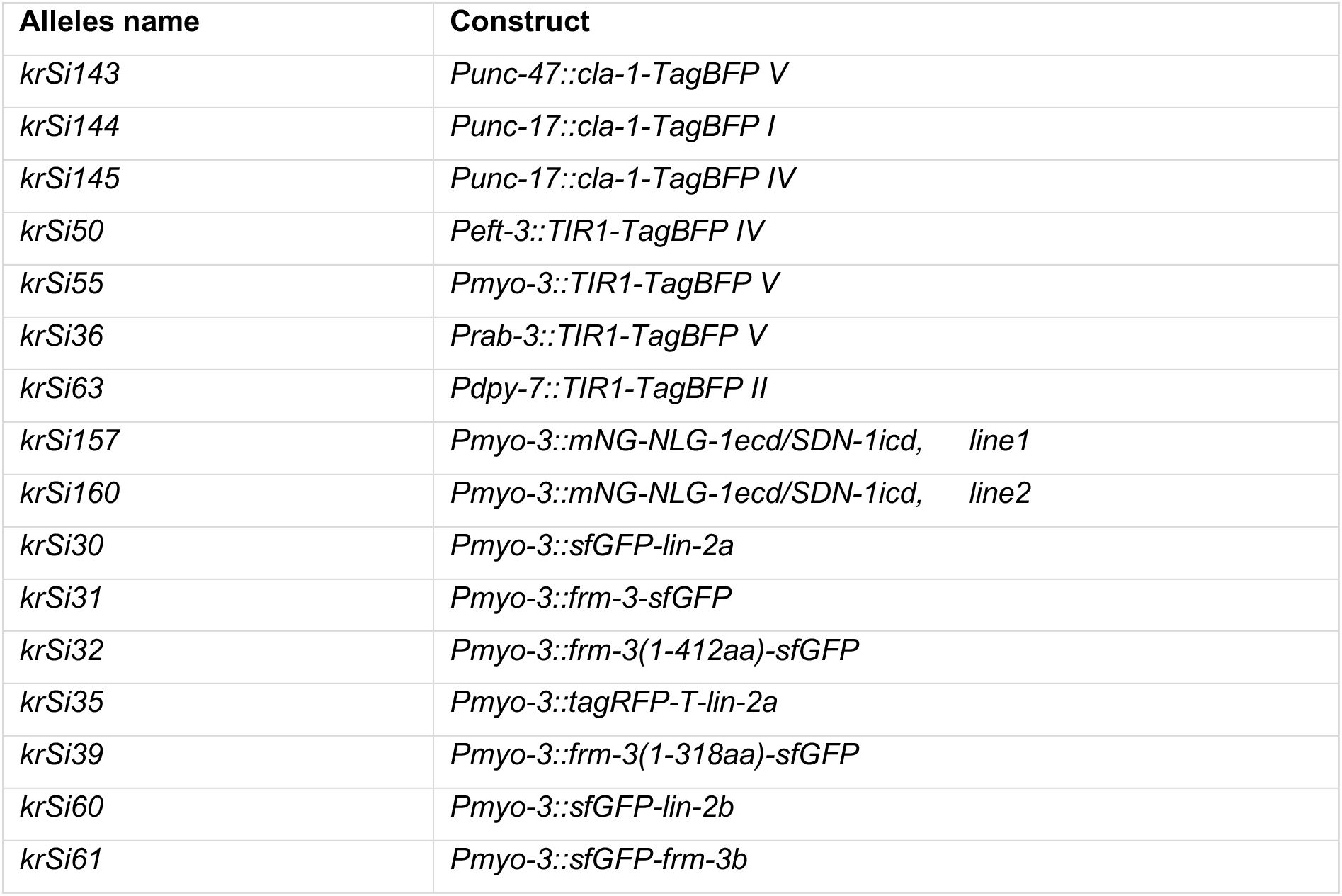
List of miniMos single-copy insertion alleles.

### Modifications of the endogenous loci

To generate the *kr383* allele, the AID sequence (Zhang et al., 2015) and mNG were inserted tandemly into the *sdn-1* locus, just before the stop codon. To generate the *kr373* and *kr387* alleles, the TagRFP-T and the wrmScarlet (Bindels et al., 2017; El Mouridi et al., 2017) sequences were inserted in the *madd-4* and in the *sdn-1* loci, respectively, just before the stop codon. The detailed protocol was described in (Dickinson et al., 2015). CrRNA sequences targeting the insertion region were designed and synthesized from IDT (Integrated DNA Technologies Inc.). CrRNA and tracerRNA (trRNA) were mixed as 1:1 ratio to form a sgRNA duplex. Adult worms were micro-injected in the gonads with a mix that contained: Cas9 nuclease 0.5 μL of 10 μg/μL (IDT Inc.), sgRNA duplex 3 μL (100 μM), repair templates (containing the DNA fragment to insert, a hygromycin resistance selection cassette surrounded by *loxP* sites, *Phsp16.41::Cre* and homology arms to target the genomic region) 50 ng/μL, pCFJ90 [*Pmyo-2::mCherry*] co-injection marker 2.5 ng/μL, RNase/DNase free water up to 10 μL. Injected animals were then grown at 25°C, and after 48h, a positive selection was performed by adding hygromycin on plates at 0.2 μg/μL. Resistant animals were then heat-shocked for 2h at 34°C by water bath to excise the hygromycin selection cassette from the genome. Worms were then screened by sequencing PCR products from the region of interest. Candidates were outcrossed once with N2 to remove unspecific background mutations.

To generate the *kr437 and kr440* alleles, PCR amplified sequences of mNG or wrmScarlet with or without homology arms were used as repair templates respectively (Dokshin et al., 2018). CrRNA was designed and synthesized by IDT (Integrated DNA Technologies Inc.). The injection mix contained annealed dsDNA donor cocktail as repair template 200 ng/μL (total 4 μg), Cas9 nuclease 0.5 μL of 10 μg /μL (IDT Inc.), sgRNA duplex 3 μL (100 μM). The candidate F1 animals were isolated by tracking the initial fluorescence knock-in. The F2 progenies were then isolated and homozygosed. The insertion was then confirmed by PCR and sequencing.

To generate *kr388*, *kr441*, *kr475*, *kr496, kr499 and kr519,* ultramer DNA repair templates were produced by IDT Inc.. The injection mix contained: Cas9 nuclease 0.5 μL of 10 μg/μL (IDT Inc.), sgRNA duplex 3 μL (100 μM), ultramer repair template 1.25 μL (100 μM), pRF4 [*Peft-3::rol-6*] co-injection marker 2.5 ng/μL, RNase/DNase free water filled up to 10 μL. Insertions in F1 roller progenies were checked by PCR. Homozygotes were isolated and the PCR product were then sent for sequencing. Candidates were outcrossed once with N2 to remove unspecific background mutations. An extensive list of modified genomic loci can be found in Table 4.

**Table 4:**
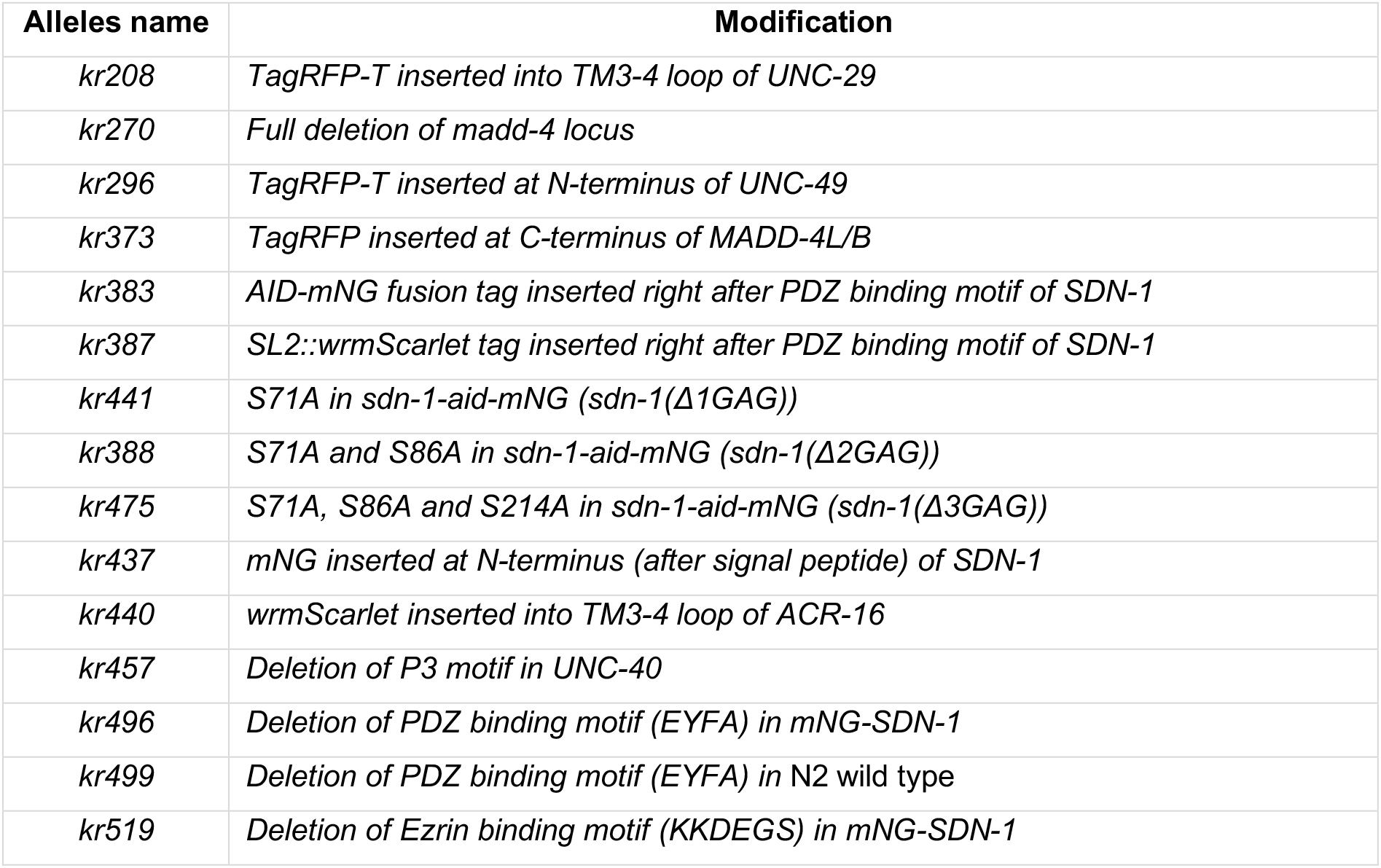
List of modified genomic loci.

| Alleles name | Modification |
| --- | --- |
| *kr208* | *TagRFP-T* inserted into TM3-4 loop of UNC-29 |
| *kr270* | Full deletion of *madd-4* locus |
| *kr296* | *TagRFP-T* inserted at N-terminus of UNC-49 |
| *kr373* | *TagRFP* inserted at C-terminus of MADD-4L/B |
| *kr383* | AID-mNG fusion tag inserted right after PDZ binding motif of SDN-1 |
| *kr387* | SL2::wrmScarlet tag inserted right after PDZ binding motif of SDN-1 |
| *kr441* | S71A in *sdn-1-aid-mNG* (*sdn-1*(Δ1GAG)) |
| *kr388* | S71A and S86A in *sdn-1-aid-mNG* (*sdn-1*(Δ2GAG)) |
| *kr475* | S71A, S86A and S214A in *sdn-1-aid-mNG* (*sdn-1*(Δ3GAG)) |
| *kr437* | mNG inserted at N-terminus (after signal peptide) of SDN-1 |
| *kr440* | wrmScarlet inserted into TM3-4 loop of ACR-16 |
| *kr457* | Deletion of P3 motif in UNC-40 |
| *kr496* | Deletion of PDZ binding motif (EYFA) in mNG-SDN-1 |
| *kr499* | Deletion of PDZ binding motif (EYFA) in N2 wild type |
| *kr519* | Deletion of Ezrin binding motif (KKDEGS) in mNG-SDN-1 |

### Auxin-induced degradation

Auxin plates were prepared by adding auxin Indole-3-Acetic Acid (Sigma) from a 400 mM stock solution in ethanol into NGM at the final concentration of 1 mM (Zhang et al., 2015). Tissue-specific degradation efficiency was monitored on animal expressing GFP fused to the Auxin-Inducible Domain (AID) under the ubiquitous promoter *Peft-3* and *Prab-3::TIR1-TagBFP*, *Peft-3::TIR1-TagBFP*, *Pdpy-7::TIR1-TagBFP* or *Pmyo-3::TIR1-TagBFP*. Animals were grown on regular or auxin plates for 3, 36, 96 and 120 hours respectively. SDN-1-AID-mNG fluorescence was scored after tissue-specific degradation on young adult animals that had hatched on regular or auxin plates.

### Microscopy imaging and quantification

For confocal imaging, young live adult hermaphrodites (24h post L4 larval stage) were mounted on 2% agarose dry pads with 5% poly-lysine beads in M9 buffer (3 g of KH_2_PO_4_, 6 g of Na_2_HPO_4_, 5 g of NaCl and 0.25 g of MgSO_4_·7 H_2_O, distilled water up to 1 L). Fluorescence images were captured using an Andor spinning disk system (Oxford Instruments) installed on a Nikon-IX86 microscope (Olympus) equipped with a 60x/NA1.2 oil immersion objective and an Evolve EMCCD camera. Each animal was imaged as a stack of optical sections (0.2 μm apart) and 41 plans were projected along Z-axis.

Quantification of images was performed using ImageJ (v1.48 by NIH) with Fiji plugin add-ons. For fluorescence intensity measurement, 50 μm (wide) × 3 μm (high) regions along the dorsal cord near the mid-body were cropped and analyzed. Acquisition settings were the same across genotypes for quantitative analysis. Each strain was imaged on at least 3 different days, and the data were pooled together. Data are presented as a percentage of the average fluorescence relative to that of the control strain. For colocalization quantification, images were captured along 40 μm of the dorsal nerve cord anterior to the vulva. The fluorescence intensity along the cord was evaluated with the Plot Profile plugin. For each channel, the values along the x-axis were normalized to the value of maximal intensity. Data are presented as min to max values for animals of each genotype. The colocalization between two channels were analyzed by Pearson’s correlation coefficient as described previously (Pinan-Lucarré et al., 2014).

To quantify the intensity of mNG-SDN-1 at ACh *vs* GABA NMJs (Fig. 1H), images of the dorsal cord of animals co-expressing *mNG-sdn-1* and *Punc-17::cla-1-BFP* or *Punc-47::cla-1-BFP* were acquired and cropped for quantification. In both channels, the background was subtracted using a rolling ball radius of 10 pixels in Fiji (Image J). The CLA-1–BFP channel of cropped images was converted to binary using the Ostu threshold. For *Punc-17::CLA-1-BFP*, noise was removed from the binary mask using the de-speckle function (twice). The thresholded image was then multiplied with the binary mask to generate the preprocessed image for further analysis. To generate the plot profiles, the intensities of each pixel along the X axis (nerve cord was converted to two-dimensional axis) were recorded for each channel in.csv files. To determine the level of mNG-SDN-1 at cholinergic synapses, the csv files were then read by an R script to define the cholinergic CLA-1-BFP domains based on values of *Punc-17::CLA-1-BFP* signals detected along the nerve cord. Next, the mean intensity of SDN-1 was calculated by measuring the area under the curve of SDN-1 profiles within cholinergic CLA-1-BFP domains and then dividing by the cholinergic CLA-1–BFP domain area. The same procedure was applied to determine the level of mNG-SDN-1 at GABAergic synapses and was also used in figure S4D to determine the level of RFP-LIN-2 level at GABA synapses. To quantify the percentage of L-ACh and GABA receptors colocalized with SDN-1 (Fig. 1G), the mNG-SDN-1 domains were defined by the same processing as described above for CLA-1-BFP. Next, the percentage of colocalized receptor was calculated by determining the ratio between the area under the curve (in receptor channel) within SDN-1 domains and the area under the curve (in receptor channel) out of the SDN-1 domains. The plot profile fluorescence signals were measured and processed by Mander’s overlapping correlation analysis modified from previous studies (Bolte and Cordelières; Dunn et al., 2011).

To quantify RFP-LIN-2 levels at GABA synapses in figures 6G, GABA domains were defined by the same processing as described for CLA-1-BFP above, using NLG-1-GFP as GABAergic synapse marker. To analyze colocalization between N-AChR and cholinergic boutons in figure 9C-D, images were processed as described in the previous paragraph.

### GST-pull down assay and heparinase treatment

GST-pull down experiment was performed based on previous protocol (Zhou et al., 2016). DNA fragments encoding the ACR-16 TM3-4 loop or the SDN-1 intracellular domain full length or lacking the EYFA PDZ binding motif were cloned to pGEX-3X expression vector and then transfected into Arctic express *E.coli* strain (Agilent). Bacterial clones were first cultured overnight in 5 mL LB medium supplemented with 100 mg/mL ampicillin. The volume of overnight-grown bacteria was enlarged in 100 mL LB/ampicillin medium by culturing 2 hours at 37 °C with shaking until OD600 reached 0.6-0.8. IPTG was added to medium at 1 mM final concentration to induce protein expression at 12 °C for 24 hours. The induced bacteria were collected by centrifugation and lysed by 20 mW pulsed sonication for 30 min in cold bacteria lysis buffer (50 mM HEPES pH 7.5, 400 mM NaCl, 1 mM DTT, 1 mM PMSF, 1 tablet of protease inhibitor cocktail/50 mL (Roche-Merck)). Protein lysate was centrifuged at 15,000 × g for 45 min at 4°C. Clear supernatants were incubated with 100 μL Glutathione sepharose-4B beads (Sigma) at 4°C for 12h. To express HA (Human influenza hemagglutinin) tagged LIN-2B, FRM-3B or FERM-FA domain, coding sequences with HA N-terminal tagged were cloned into pcDNA3.1(+) expression vector. Fusion proteins were expressed by TnT® Quick Coupled system (Promega). 15 μL of HA tagged protein was then incubated with candidate GST-coated beads at 4°C for 2 hours. The beads were washed five times with STE (10 mM Tris-Cl pH8.0, 150 mM NaCl, 1 mM EDTA, 0.1% Tween-20) buffer. Proteins were eluted by adding equal volume of 2xSDS sampling buffer (Bio-rad) and boiled for 10 min. Samples were electrophoresed in 12% SDS/PAGE (Bio-rad). The separated proteins were incubated with an anti-HA antibody (#3724, Cell signaling Tech.) at 1:1000 dilution. The GST fusion loading amount was detected by stain free gel imaging or Ponceau S red staining immediately after membrane transfer.

To digest heparan sulfate chains of SDN-1 and to compare with GAG binding site mutant animals, mNG-SDN-1 non-mutated or *Δ*1, 2 and 3 GAGs fusion proteins were first concentrated by immunoprecipitation of roughly 20,000 nematodes per genotype using Trap-A bead coupled with anti-mNG nanobody (Chromotek). 1.5 U of heparanase I and III (Sigma) was added to non-mutated tube containing 500 μL washing buffer (50 mM HEPES, pH 7.7, 50 mM NaCl) and 100 Trap-A beads. The tube was then incubated at 37°C for 2 hours. The proteins of each group were eluted by adding 100 μL 2x sampling buffer and boiled for 10 min. Samples were separated by 12% SDS/PAGE (Bio-rad) and blotted with anti-mNG antibody at 1:1000 dilution. N2 wild-type animals were used as negative control.

### Electrophysiology

Adult nematodes were glued (Histoacryl Blue, B. Braun) along the dorsal side of the body to the surface of a plastic culture dish (Corning). A sharpened tungsten rod (A-M systems Inc) was used to perform a lateral incision before the vulva and to remove the viscera. The cuticle flap was glued back to expose the ventral body-wall muscles and the preparation was treated by collagenase type IV for 20 s at the concentration of 0.5 mg/mL. Recordings were performed on the two first ventromedial muscle cells before the vulva from the right muscle quadrant. For all recordings, strains express ChR2 in cholinergic motoneurons (*zxIs6*) (Liewald et al., 2008). Animals were grown on NGM plates freshly seeded with 300 μL of OP50 culture supplemented by 0.25 μL of 100 mM all-*trans* retinal stock (dissolved in ethanol). Membrane currents were recorded in the whole-cell configuration using a MultiClamp 700B amplifier (Molecular Devices). Acquisition and command voltage were done using the Clampex 10 software driving an Axon Digidata 1550 (Molecular Devices). The resistance of recording pipettes ranged between 3 and 4 MΩ. Data were analyzed with Clampfit 10 (Molecular Devices) and graphed with Origin software (OriginLab).The bath solution contained (in mM) 150 NaCl, 5 KCl, 5 CaCl_2_, 1 MgCl_2_, 10 glucose, 15 HEPES, and sucrose to 320 mOsm/L (pH 7.2), and the pipette solution (in mM) 120 KCl, 4 NaCl, 5 EGTA, 10 TES, 4 MgATP and sucrose to 310 mOsm/L (pH 7.2).

For currents evoked by cholinergic motoneuron stimulation, the slit worm preparations were exposed to 10 ms light stimulation performed with the pE2 system (CoolLED) at a wavelength of 460 nm drived by Clampex 10 software. The protocol was the following: a first stimulation triggered the activation of L-AChR and N-AChR; the peak current value was measured and named total peak current. The preparation was then perfused with bath solution supplemented with dihydro-beta-erythroidine (DHβE) at the concentration of 0.01 mM leading to the specific inhibition of N-AChR. In these conditions, a second stimulation was performed activating only L-AChR. A third stimulation was then applied to check if the current was stable and the value of the peak current was measured and named L-AChR evoked current. Finally, for each cell, the L-AChR evoked current was subtracted from the total peak current to obtain the N-AChR evoked current values.

For agonist-evoked currents, nicotine and levamisole diluted at the concentration of 0.1 mM in the bath solution were pressured-ejected during 100 ms from a pipette similar in size to the patch pipette at 10 psi using a PDES-2DX system (NPI electronic). The slit worm preparations were continuously perfused with fresh bath solution via a gravity flow delivery system. All chemicals were obtained from Sigma-Aldrich except DHβE provided by Tocris. All experiments were done at 20°C.

## SUPPLEMENTAL INFORMATION

**Figure S1.**
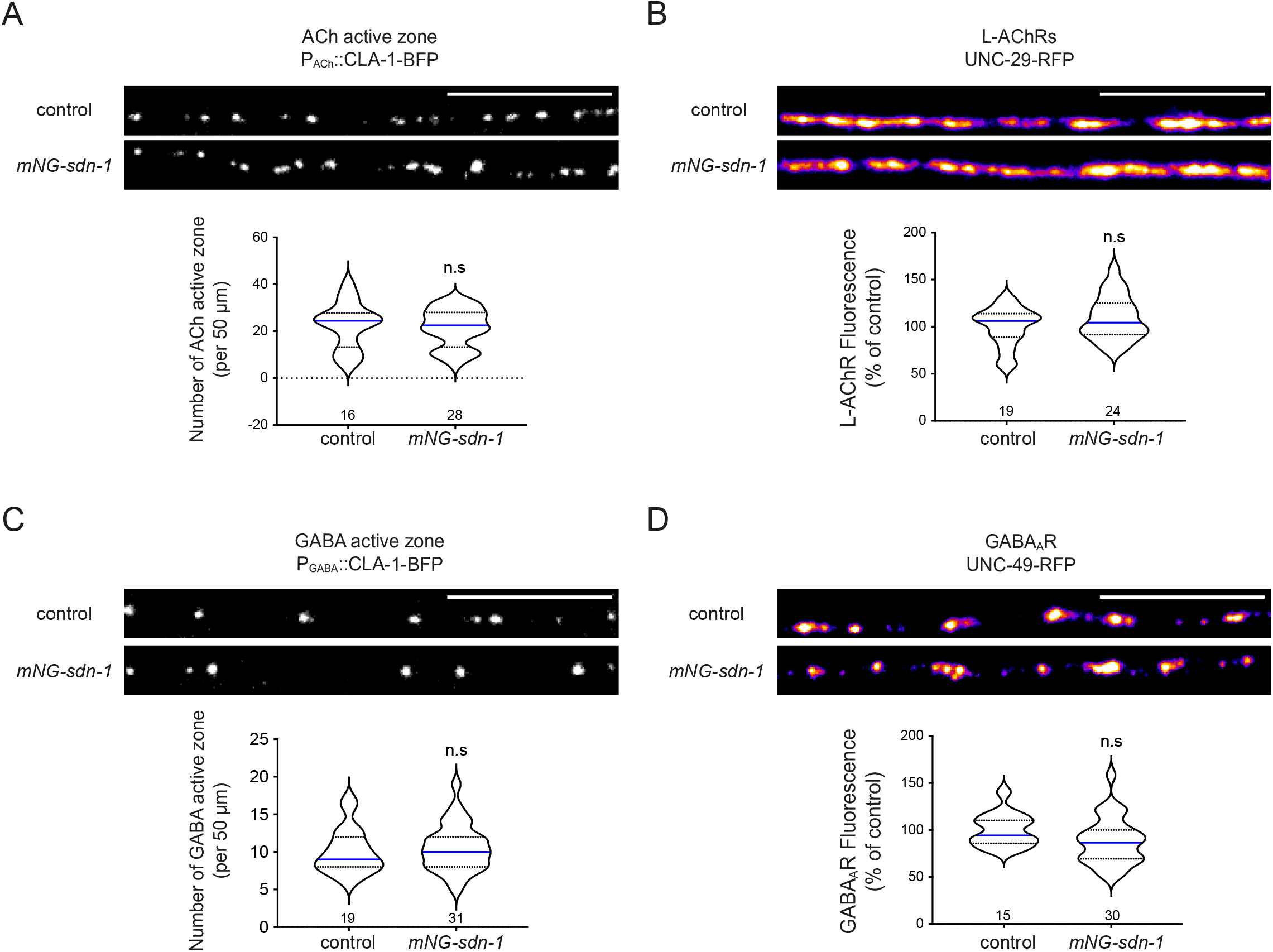
mNG insertion in the *sdn-1* locus has no effect on NMJ organization. (A) Confocal detection of the presynaptic active zone marker CLA-1-BFP under the control of the cholinergic neuron-specific promoter *Punc-17* in control and *mNG-sdn-1* knock-in animals. Bottom: quantification of the number of active zones per 50 μm of dorsal cord in control and *mNG-sdn-1* animals. (B) Confocal detection and quantification of RFP-labeled L-AChR in control and *mNG-sdn-1* animals. Fluorescence levels were normalized to the mean value of control. Mann-Whitney test, n.s: not significant. (C) Confocal detection of the presynaptic active zone marker CLA-1-BFP under the control of the GABAergic neuron-specific promoter *Punc-47* in control and *mNG-sdn-1* animals. Bottom: quantification of the number of active zones per 50 μm of dorsal cord in control and *mNG-sdn-1* worms. (D) Confocal detection and quantification of RFP-labeled GABA_A_R in control and *mNG-sdn-1* animals. Fluorescence levels were normalized to the mean value of control. Scale bars = 10 μm. Violin plots as in figure 1G. Mann-Whitney tests, n.s: not significant.

**Figure S2.**
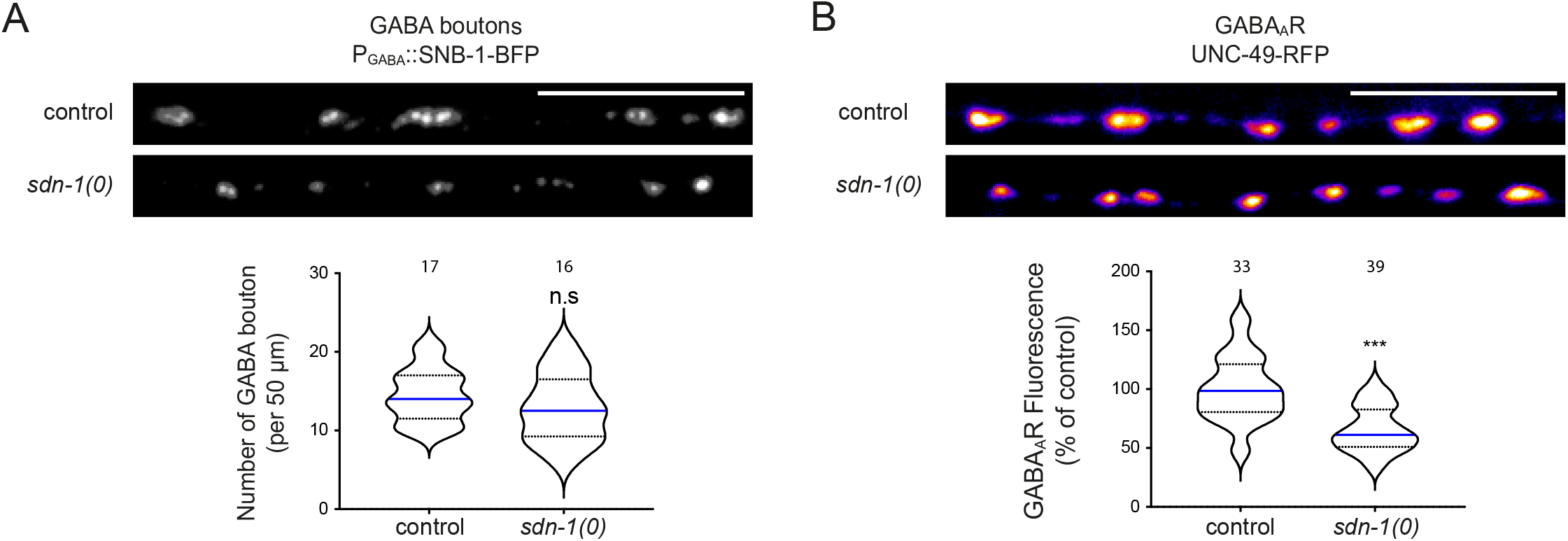
SDN-1 has a moderate effect on GABA_A_R synaptic content. (A) Confocal detection of the presynaptic active zone marker CLA-1-BFP under the control of the GABAergic neuron-specific promoter *Punc-47* in control and *sdn-1(0)* mutant animals. Bottom: quantification of the number of active zones per 50 μm of dorsal cord in control and *sdn-1(0)* worms. (B) Confocal detection and quantification of RFP-labeled GABA_A_R in control and *sdn-1(0)* animals. Fluorescence levels were normalized to the mean value of control. Scale bars = 10 μm. Violin plots as in figure 1G. Mann-Whitney tests, ***p<0.0001, n.s: not significant.

**Figure S3.**
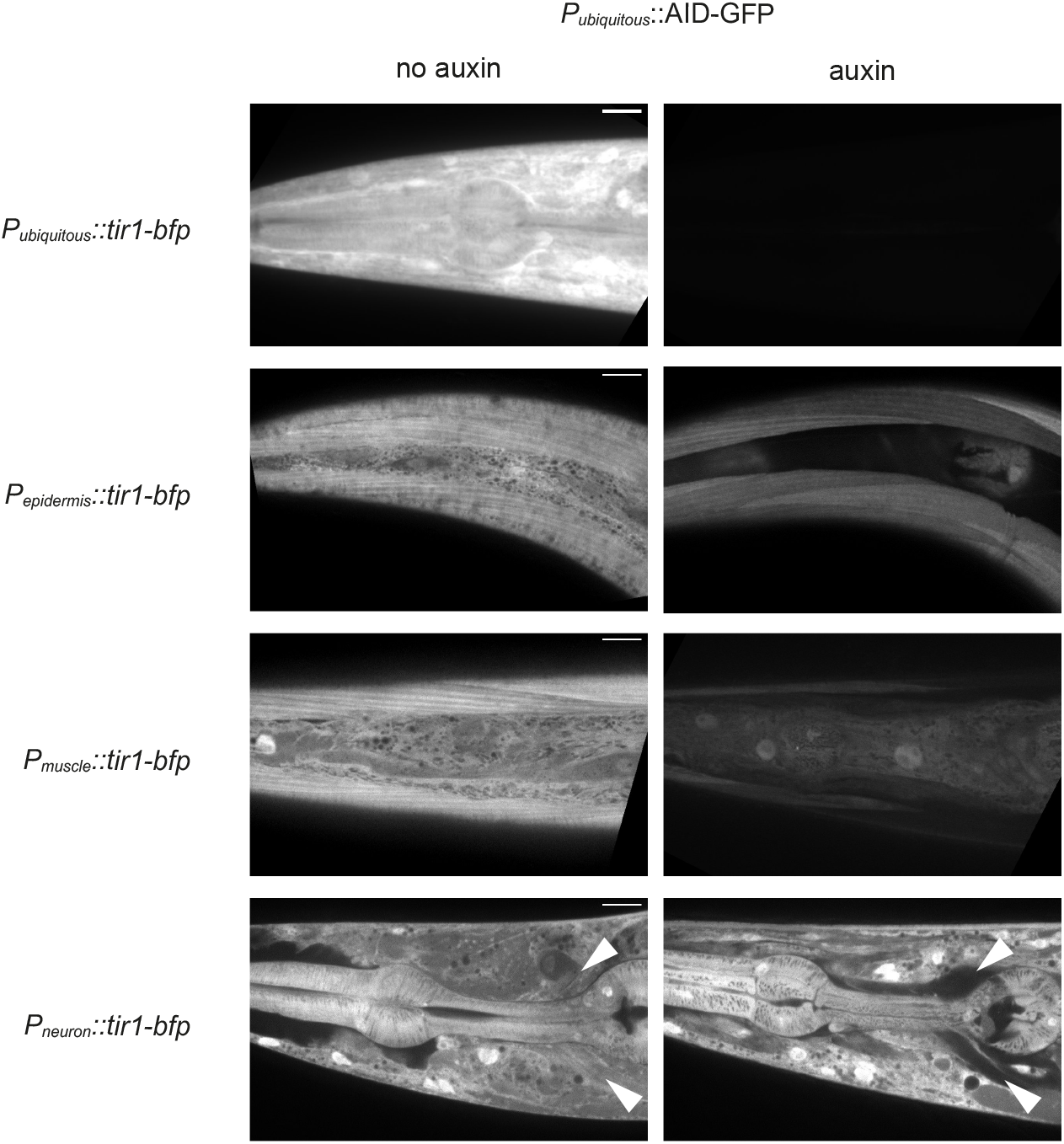
The Auxin-Inducible-Degron (AID) leads to tissue-specific degradation. Confocal detection of AID-GFP expressed in all tissues by the ubiquitous promoter *Peft-3* from animals grown on regular NGM plates (control, left panels) or on auxin plates (right panels). Tissue-specific degradation was achieved by co-expressing AID-GFP and *Peft-3::tir-1-bfp* (ubiquitous)*, Pdpy-7::tir-1-bfp* (epidermis)*, Pmyo-3::tir-1-bfp* (body-wall muscle), or *Prab-3::tir-1-bfp* (neurons). White arrowheads indicate the position of the nerve ring. Scale bars = 10 μm.

**Figure S4.**
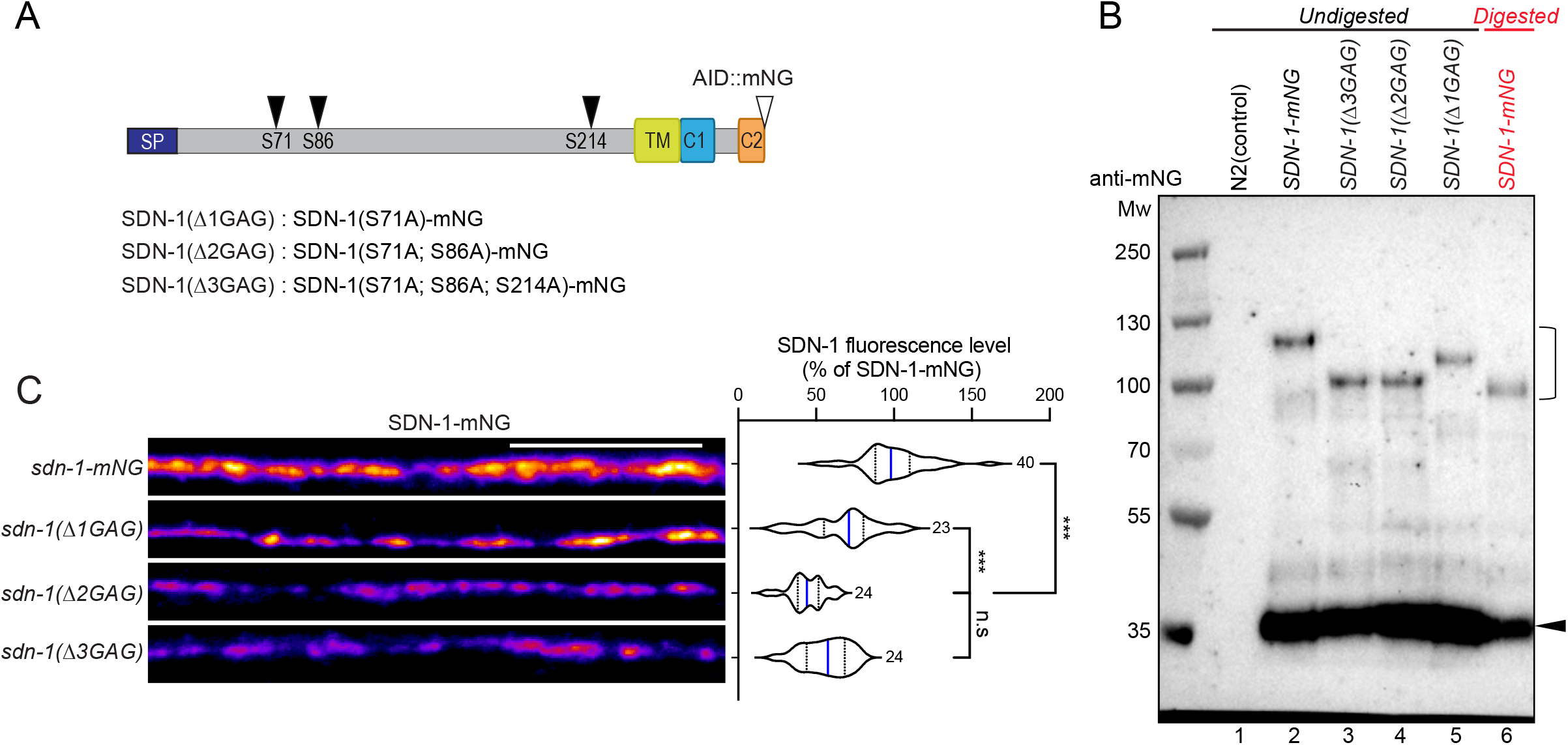
Heparan sulfate chains modulate SDN-1 synaptic content but are dispensable for its MADD-4-dependent synaptic localization. (A) Protein structure of SDN-1-mNG with the position of GAG chain anchoring serines. Ser71, Ser86 or Ser214 were mutated into Alanine in mutants lacking one (Δ1GAG), two (Δ2GAG) or three (Δ3GAG) GAG chain anchoring sites. Detailed domain information as in Fig 1A. (B) Western blot showing the migration of non-mutated SDN-1-mNG (lane 2), SDN-1(Δ3GAG)-mNG (lane 3), SDN-1(Δ2GAG)-mNG (lane 4), SDN-1(Δ1GAG)-mNG (lane 5) protein precipitated from worm lysate. Expected SDN-1-mNG molecular weight is 58 kDa. Bracket likely indicates the SDS resistant dimerization form of SDN-1. Black arrowhead indicates potential cleaved C-terminus-mNG band. Protein samples from lanes 1-5 were undigested while sample from lane 6 (shown in red) was digested by Heparinase I and III for 2 hours at 37 °C. Protein lysate from N2 wild-type animals was used as control in lane 1. (C) Confocal detection of SDN-1-mNG with no modification or CRISPR mutation of one (Δ1GAG), two (Δ2GAG) or three (Δ3GAG) GAG chain anchoring sites as specified in A. Fluorescence levels were normalized to the mean of non-mutated SDN-1-mNG fluorescence. One-way ANOVA followed by Tukey’s multiple comparison tests. ***p<0.001, n.s: not significant. Violin plot as in figure 1G. Scale bars = 10 μm.

**Figure S5.**
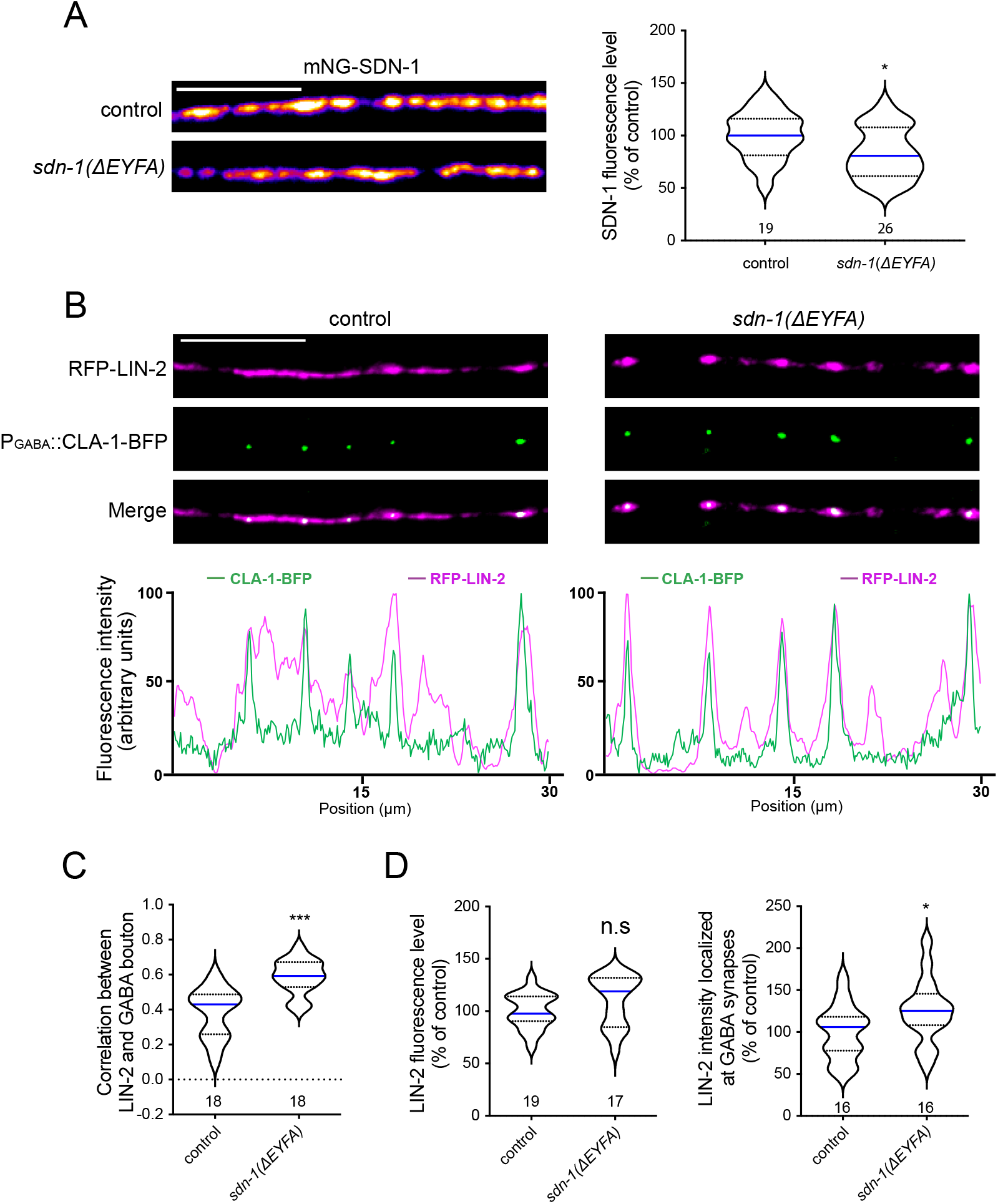
LIN-2 is concentrated at GABA synapses in animals lacking SDN-1 PDZ binding motif. (A) Confocal detection and quantification of mNG-SDN-1 fluorescence levels in control (*mNG-sdn-1*) and *mNG-sdn-1*(*ΔEYFA*) animals. Fluorescence levels were normalized to the mean value of control animals. (B) Confocal detection of RFP-LIN-2B and the presynaptic active zone marker CLA-1-BFP at GABA synapses in control and *sdn-1(ΔEYFA)* animals expressing *Pmyo-3::rfp-lin-2b* and *Punc-47::cla-1-bfp*. The fluorescence profiles indicate muscle-expressed RFP-LIN-2B and GABA synaptic CLA-1-BFP fluorescence intensities along the nerve cord from the pictures above. (C) Pearson’s correlation coefficient between RFP-LIN-2B and the GABAergic presynaptic active zone marker CLA-1-BFP in control and *sdn-1(ΔEYFA)* animals. (D) Fluorescence levels of RFP-LIN-2B at the dorsal nerve cord or only at GABA synapses of control and *sdn-1(ΔEYFA)* animals. Total fluorescence (left panel) was measured as for other figures. To measure RFP-LIN-2B fluorescence specifically localized at GABA synapses (right panel), GABA synaptic CLA-1-BFP marker was used to define GABA regions from processed images and RFP-LIN-2B fluorescence was quantified in these defined regions (see material and methods for details). Fluorescence levels were quantified and normalized to the control groups. Scale bars = 10 μm. Violin plots as in figure 1G. Mann-Whitney tests, ***p<0.001, *p<0.05, n.s: not significant.

## Notes

### Competing Interest Statement

The authors have declared no competing interest.

